# Timing of standard chow exposure determines the variability of mouse phenotypic outcomes and gut microbiota profile

**DOI:** 10.1101/2024.03.28.587032

**Authors:** Megan M. Knuth, Carolina Vieira Campos, Kirsten Smith, Elizabeth K. Hutchins, Shantae Lewis, Mary York, Lyndon M. Coghill, Craig Franklin, Amanda MacFarlane, Aaron C. Ericsson, Terry Magnuson, Folami Ideraabdullah

## Abstract

Standard chow diet contributes to reproducibility in animal model experiments since chows differ in nutrient composition, which can independently influence phenotypes. However, there is little evidence of the role of timing in the extent of variability caused by chow exposure. Here, we measured the impact of diet (5V5M, 5V0G, 2920X, and 5058) and timing of exposure (adult exposure (AE), lifetime exposure (LE), and developmental exposure (DE)) on growth & development, metabolic health indicators, and gut bacterial microbiota profiles across genetically identical C57BL6/J mice. Diet drove differences in macro-and micronutrient intake for all exposure models. AE had no effect on measured outcomes. However, LE mice exhibited significant sex-dependent diet effects on growth, body weight, and body composition. LE effects were mostly absent in the DE model, where mice were exposed to chow differences from conception to weaning. Both AE and LE models exhibited similar diet-driven beta diversity profiles for the gut bacterial microbiota, with 5058 diet driving the most distinct profile. Diet-induced beta diversity profiles were sex-dependent for LE mice. Compared to AE, LE drove 9X more diet-driven differentially abundant genera, majority of which were the result of inverse effects of 2920X and 5058. Our findings demonstrate that lifetime exposure to different chow diets has the greatest impact on reproducibility of experimental measures that are common components of preclinical mouse model studies. Importantly, weaning DE mice onto a uniform diet is likely an effective way to reduce unwanted phenotypic variability among experimental models.

## Introduction

Standard rodent (chow) diets contain adequate nutrition but vary greatly in macro-and micronutrient composition^1^. Sources of nutrients vary from diet to diet, and complete ingredient lists are not always provided^1,2^. Since differences in nutrient intake and other naturally occurring bioactive compounds can independently influence the phenotypic outcome^3,4,5,6^, the use of different chows between labs or model development steps is a major problem for the reproducibility of research findings. This is especially problematic for studying preclinical animal models of disease, which often require accurate repeated measures of the isolated phenotypic effects of genotype, surgery, or pharmaceutical drug exposure. For example, in mice with acetaminophen-induced liver injury, a chow diet determined the extent of liver damage^7^. In another mouse study, standard chow determined host immunity response to influenza^8^.

Previous studies also show how chow diet differences alone can drive significant differences in mouse model phenotypes. A hexokinase II knockdown study demonstrated significant effects of chow on mouse body weight, heart weight, and hexokinase phenotypes^9^. Another example comparing the effects of standard chow to a purified diet (AIN-76A) found that the chow diet drove differences in food intake and plasma essential amino acid levels in inbred C57BL/6 mice^10^. Lastly, differences in chow diets with variable fiber sources were found to induce differences in the distribution of gut bacterial communities in outbred CD-1 mice^11^.

Epigenetic changes are often implicated as mechanistic drivers of the phenotypic variability caused by diet. Although this has not been extensively studied for chow diets, several studies using mouse models demonstrate the effects of isolated nutrients. For example, a high-fat diet (HFD) induced histone modifications in mouse adipose tissue that were then linked to metabolic syndrome-like phenotypes^12^. A separate study showed that HFD induced epigenetic modifications in mouse brain that caused learning and memory deficits^13^.

Most studies on the effects of the chow diet have focused on adult exposures, but research in animal models and human populations shows that developmental windows are particularly susceptible to dietary changes^14^. Data demonstrating the effect of diet timing on outcome within the same study are limited, but a recent study in Swiss Webster mice found that postnatal exposure to HFD at different stages of life had differing effects on diabetic outcomes^15^. HFD exposure across the lifespan (postnatal day (PND) 2-325) resulted in the greatest impact, while developmental exposure between birth and weaning (PND 2-21) had the second greatest impact^15^. Surprisingly, slightly longer developmental exposure, including pre-and postweaning (PND 2-35), had the least impact^15^.

Here, we measured the phenotypic impacts of exposing mice at different life stages to commonly used standard chows (5V5M, 5V0G, 2920X, and 5058). Three different exposure windows were compared using a multigenerational parallel treatment scheme with inbred mice to limit effects to those within a single genetically and environmentally controlled population of mice: (1) Adult exposure (AE) – parents, (2) Lifetime exposure (LE) – offspring, and (3) Developmental exposure (DE) – siblings of LE offspring. Females and males were tested to address sex effects on the most common preclinical mouse model strain background, C57BL6/J. Differences in the macro-and micronutrient composition of chow diets were measured, and the impacts on nutrient intake, growth and development, metabolic health indicators, and gut bacterial microbiota composition were assessed.

## Results

### Measured values for micronutrients are variable among standard chow diets and far exceed the recommended levels

To assess differences in nutrient exposure from the chow diets, we measured nutrient levels in the four chows fed to mice in this study (**Supplemental Table 1**). We compared these to the manufacturer’s estimates on the diet spec sheets (**Supplemental Table 2**). Macronutrient composition and the levels of two micronutrients with important roles in development (vitamin D & folic acid) were assessed. We found that total measured calories, carbohydrate, fat, and protein content were relatively similar among the four diets except for 2920X, which contained the lowest total calories and fat and the highest carbohydrate content (**Figure 1a-b**). This was consistent with the manufacturer’s estimates (**Figure 1a-b**). Measured fiber content was variable among the diets with 2920X containing the highest total and soluble fiber and 5V0G containing the highest crude fiber (both ∼1.3× higher than the lowest, 5V5M) (**Figure 1c**). Only crude fiber levels were provided in the manufacturer estimate; this pattern was similar (**Figure 1c**).

**Figure 1.**
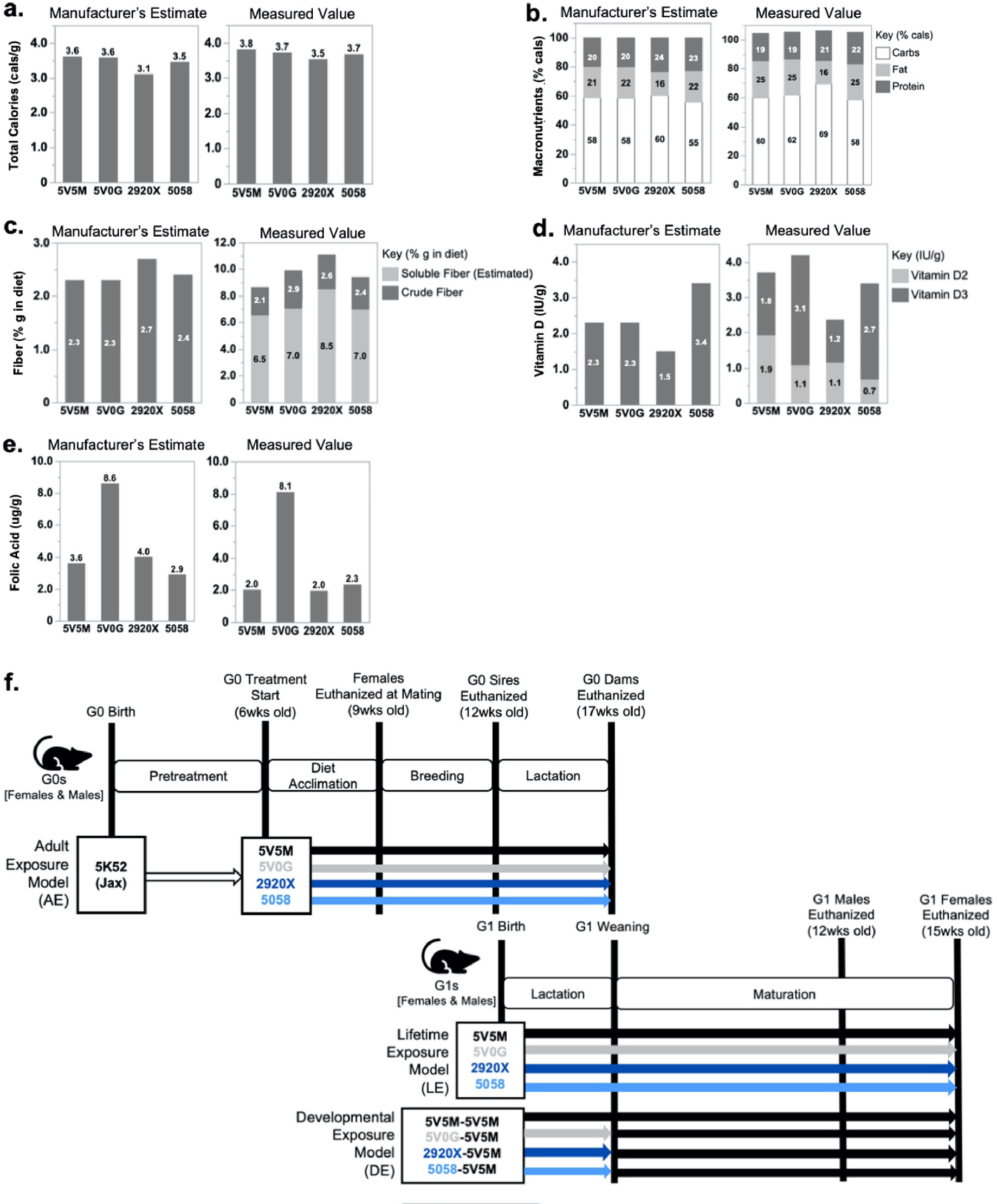
Standard chow diet composition and dietary treatment scheme. **a-e.** The macro-and micronutrient estimates provided by the manufacturer are compared to measured values for **a.** total calories; **b.** macronutrients (fat, carbs, protein); **c.** fiber (only crude fiber content was provided by the manufacturer); **d.** vitamin D (only vitamin D3 levels were provided by the manufacturer); and **e.** folic acid. **f.** Standard chow dietary treatment scheme for adult exposure (AE), lifetime exposure (LE), and developmental exposure (DE) models. AE model: C57BL/6J mice at 6 weeks of age were equally divided across four standard chow models (5V5M, 5V0G, 2920X, and 5058) for 3 weeks of chow diet acclimation. After 3 weeks, a subset of AE females was euthanized for micronutrient measures. The remaining AE females and males were mated in breeder trios based on chow diet assignment. AE males were euthanized after 3 weeks of mating, and females were euthanized at weaning. LE & DE models: At weaning pups were caged by litter and sex, and each litter was divided between LE and DE. LE mice were weaned onto the diet assignment of their parents, while DE mice were weaned onto the 5V5M diet and remained until the end of the study.

Vitamin D and folic acid concentrations were more variable among the diets and less consistent with the manufacturer’s estimates, which are based on added micronutrients and “do not account for contributions from other ingredients”. For vitamin D, 1 IU/g of feed is considered “*adequate*” and potentially even a *“considerable excess*” for laboratory mouse chow^16^. Total vitamin D content (D2 + D3) measured for our standard chows ranged from 2.3 IU/g (2920X) to 4.2 IU/g (5V0G). This was considerably higher (∼2-4×) than the recommended amounts and higher than the manufacturer’s estimates of 1.5 IU/g (2920X) to 3.4 IU/g (5058) (**Figure 1d**). Vitamin D2 contributed to this outcome since the manufacturer’s estimate only provided vitamin D3 values. However, our measured vitamin D3 values alone differed from the manufacturer’s estimate. (**Figure 1d**). The recommended dietary folic acid intake for mice is 2 ug/g of diet^17^. Measured folic acid values were lower than the manufacturer’s estimates and consistent with the recommended value (∼2-2.3 ug/g), except for 5V0G. 5V0G measured folic acid level was similar to the manufacturer’s value and 4× higher than the other diets and the recommended value, and more similar to a folic acid-supplemented diet (10 mg/kg^18^) (**Figure 1e**).

### Standard chow diets impact nutrient exposure differently depending on the diet and timing of exposure

We first assessed the effect of adult exposure (AE) to different standard chow diets. C57BL/6J females and males were fed one of four standard chows that differed in nutrient composition (**Figure 1f**, **Supplemental Tables 1-2**). After 3 weeks of acclimation to the diet, females were randomly split into two groups: Group 1 was euthanized to measure nutrient status after diet acclimation but before mating, and Group 2 remained on the diet while mating with males to measure nutrient status and phenotypic outcomes after breeding (**Figure 1f**).

Nutrient exposure is determined in part by nutrient intake, which is a combination of dietary content and food intake. Food intake in the AE model was relatively similar among the diets except for 5058, which had ∼1.6× higher food intake than the lowest (5V5M) in females before mating (**Figure 2a-b**, **Supplemental Figure 1a-b**). Less variability was observed in the males before mating and during breeding when the males and females were combined (**Figure 2a-b**, **Supplemental Figure 1a-b**). Intake of total calories, macronutrients, and fiber mirrored the food intake variability among the diets, with 5058 exhibiting the highest nutrient intake for females before mating (**Figure 2c-e**). In contrast, micronutrient intake more closely mirrored the diet composition (**Figure 2f-g**). Vitamin D intake was the most variable in females before mating, with ∼2.3× difference between the highest (5058) and lowest (2920X) (**Figure 2f**). Folic acid intake was highly variable before and after breeding, with a ∼4× difference between the highest (5V0G) and lowest (5V5M) (**Figure 2g**). These differences in intake did not drive significant differences in serum vitamin D (25(OH)D) or folate concentrations (**Supplemental Figure 2**).

**Figure 2.**
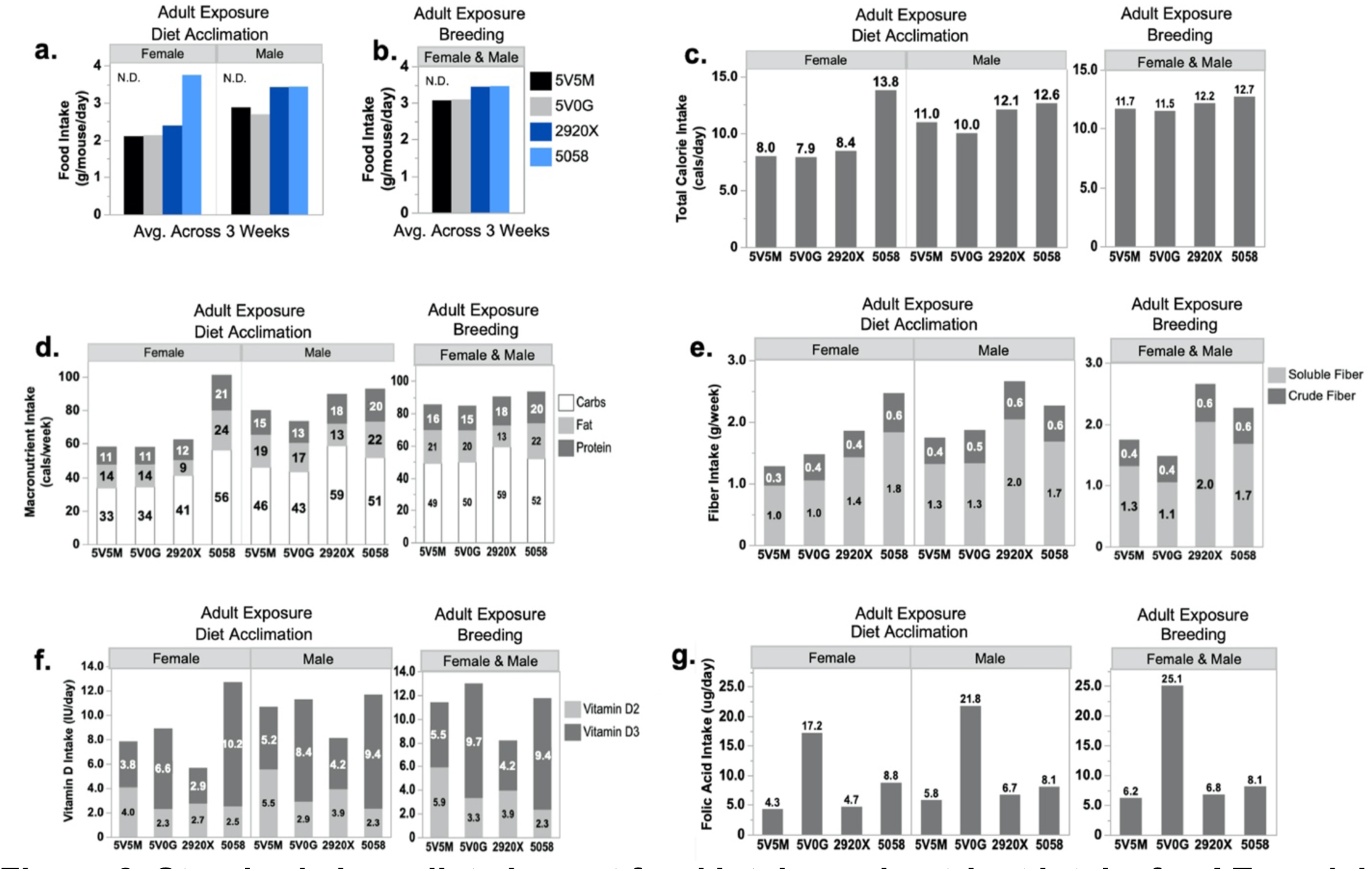
Standard chow diets impact food intake and nutrient intake for AE model. Food intake was measured weekly for each cage, and average daily values were calculated as g/mouse/day for females and males during **a.** diet acclimation and **b.** breeding. Average nutrient intake was calculated based on food intake, measured macro-and micronutrient values, and reported for **c**. total calories; **d.** macronutrients (fat, carbs, protein); **e.** fiber; **f.** vitamin D; and **g.** folic acid. Diet effects were calculated by one-way ANOVA or Kruskal-Wallis test and *significant when p<0.05. N.D. = p-value not determined due to small sample size.

Offspring generated from breeders in the AE model was used to assess the effects of developmental exposure (DE) and lifetime exposure (LE, development + adult) to different chow diets. LE mice were exposed to one of four standard chow diets from conception to adulthood (**Figure 1f**). DE mice were siblings of LE mice that were only exposed to different diets from conception to weaning and then placed on a uniform diet until adulthood (**Figure 1f**).

Consistent with findings in the AE model, food intake in the LE model significantly differed among the diets, with 5058 mice eating the most and 5V5M the least (**Figure 3a**, **Supplemental Figure 3a**). However, in contrast with the AE model, LE males and females were both affected. Less variability in food intake among the diets was observed in the DE model, although 5058 remained the diet with the highest intake, and 2920X intake was seemingly reduced (**Figure 3b**, **Supplemental Figure 3b**). The variability in nutrient intake among the diets in the LE model was also very similar to the AE model, except both males and females were affected (**Figure 3c-g**). Intake of both macro-and micronutrients in the DE model (where mice were fed the same 5V5M diet from weaning to adulthood) closely mirrored the patterns of food intake with the highest nutrient intakes for 5058-5V5M and lowest for 2920X-5V5M (**Figure 3c-g**). In contrast with the AE model, chow diet differences in the LE model drove significantly different serum vitamin D (25(OH)D) concentrations (**Supplemental Figure 4a**). However, serum 25(OH)D differences did not reflect vitamin D intake (**Figure 3f**). Instead, 2920X, which had the lowest vitamin D intake, exhibited the highest mean serum 25(OH)D levels and 5V5M exhibited the lowest (**Supplemental Figure 4a**). Chow diets did not drive significant differences in serum vitamin D for the DE model nor folate concentrations for the LE or DE model (**Supplemental Figure 4b-d**).

**Figure 3.**
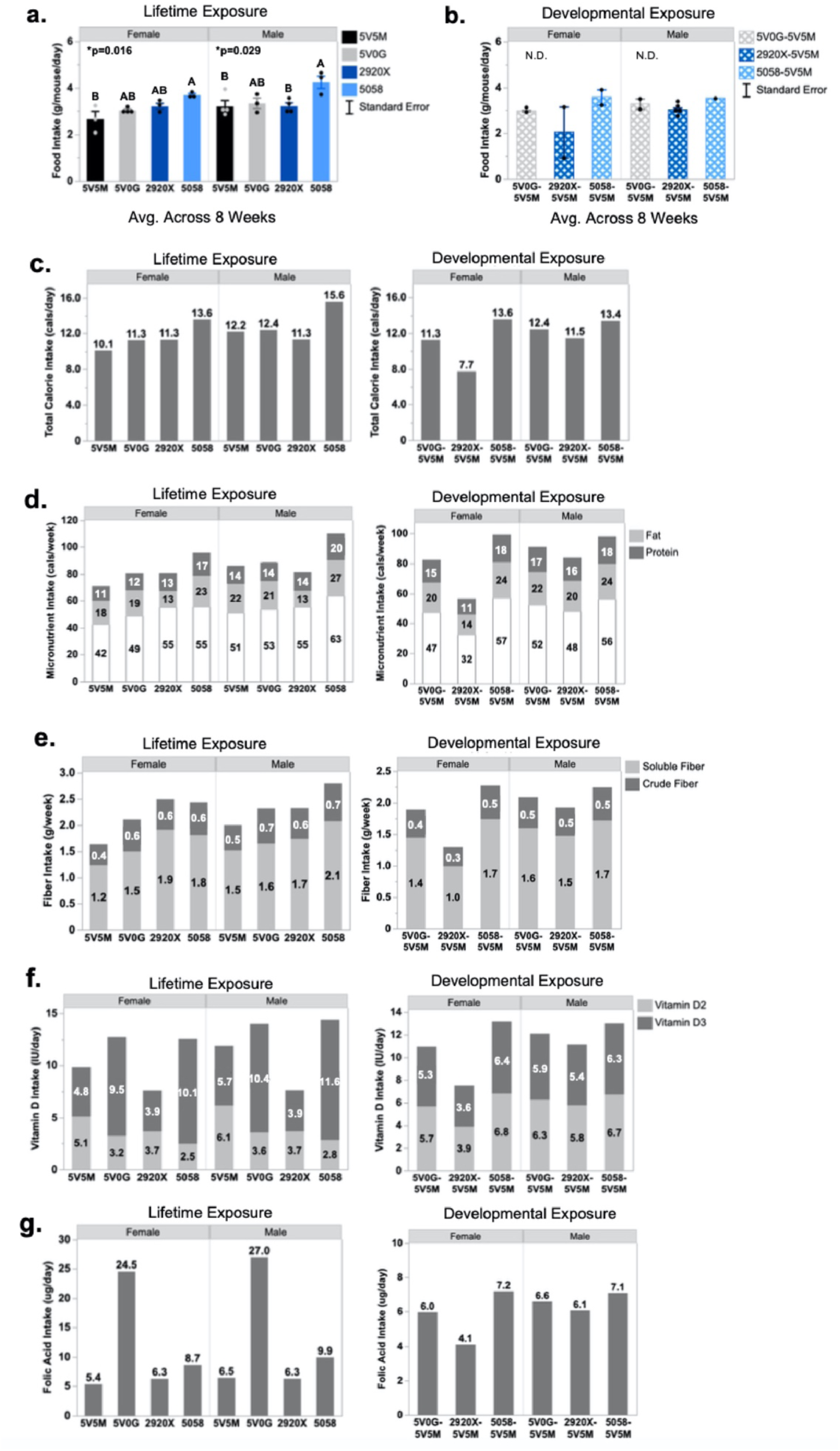
Standard chow diets impact food intake and nutrient intake for LE and DE models post-weaning. Food intake was measured weekly for each cage, and average daily values were calculated as g/mouse/day for females and males for **a.** LE and **b.** DE models. Average nutrient intake was calculated based on food intake, measured macro-and micronutrient values, and reported for **c.** total calories; **d.** macronutrients (fat, carbs, protein); **e.** fiber; **f.** vitamin D; and **g.** folate. LE and DE models were analyzed separately. Error bars represent the standard error of the mean for all graphs. Diet effects were calculated by one-way ANOVA or Kruskal-Wallis test and *significant when p<0.05. N.D. = p-value not determined due to small sample size.

### Lifetime exposure to different standard chow diets drives the greatest variability in offspring growth and adiposity

Given the substantial differences in macro-and micronutrient intake among the four diets tested, we assessed effects on phenotypes in the AE, LE, and DE models. Despite the large differences in nutrient intake among chow diets in the AE model, up to 11 weeks of adult chow diet exposure had no significant effect on metabolic health indicators in females, including body weight (**Figure 4a**, **Supplemental Figure 5a**), perigonadal white adipose tissue (PWAT) weight (**Figure 4b**), or fasting blood glucose (**Figure 4c**). Males were similarly unaffected (**Figure 4d-f**, **Supplemental Figure 5b**), although statistical testing was not possible due to the small sample size (n = 2/diet). The AE model also showed no significant diet effects on breeding success metrics (**Table 1**). Thus, adult exposure to these chow diets during breeding likely has minimal effects on phenotypes similar to those measured here.

**Figure 4.**
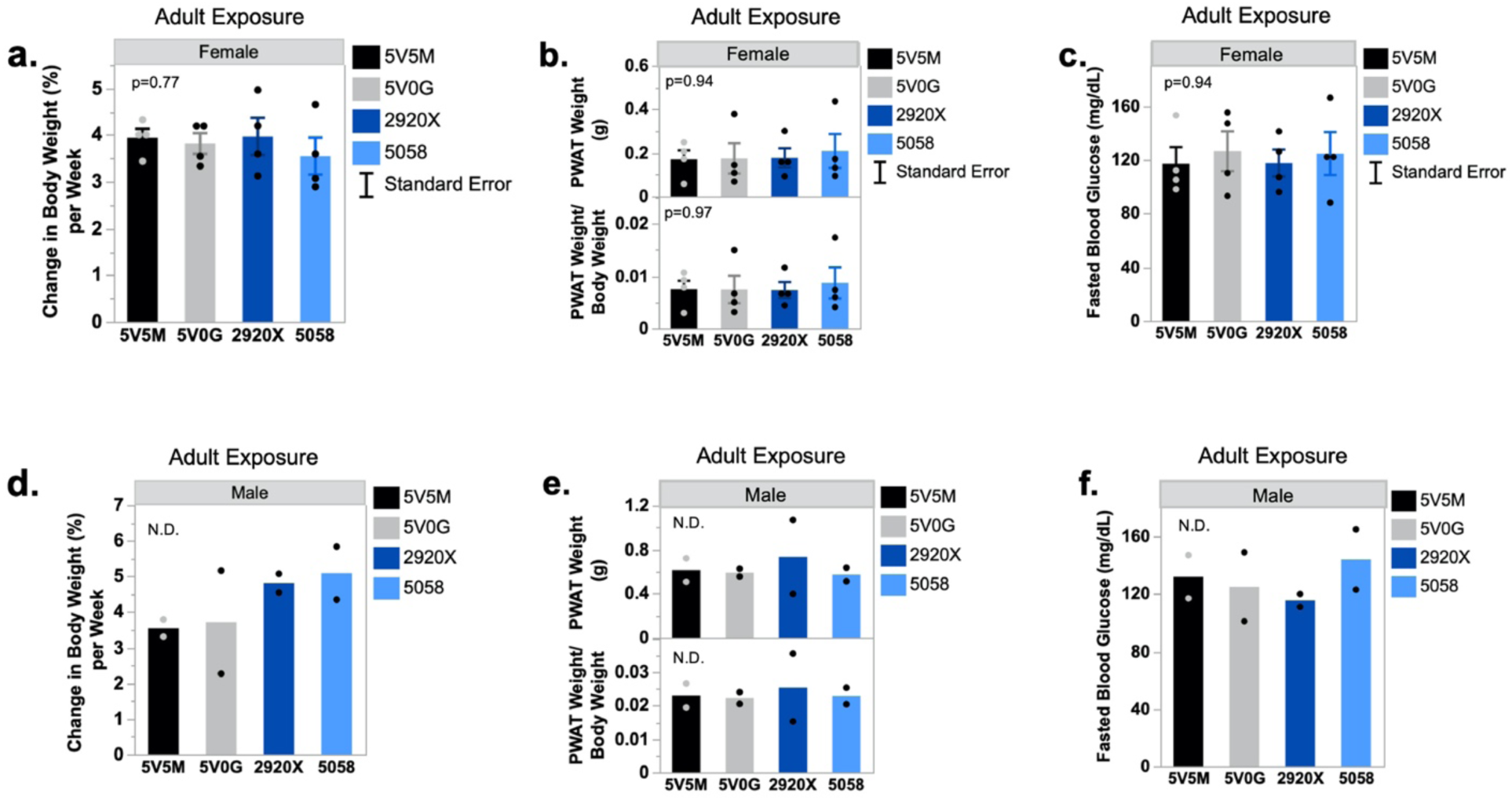
No effect of standard chow diets on AE model phenotypes. **a.** Change in female body weight from diet start to diet end. **b.** Female perigonadal white adipose tissue (PWAT) weight (g) and PWAT weight (g) relative to body weight (g) collected after euthanasia. **c.** Fasting blood glucose was measured in live females following a 12-hr fast. **d-f.** The same measures were assessed in male mice. Error bars represent the standard error of the mean for all graphs. Diet effects were calculated by one-way ANOVA or Kruskal-Wallis test and *significant when p<0.05. N.D. = p-value not determined due to small sample size.

**Table 1.**
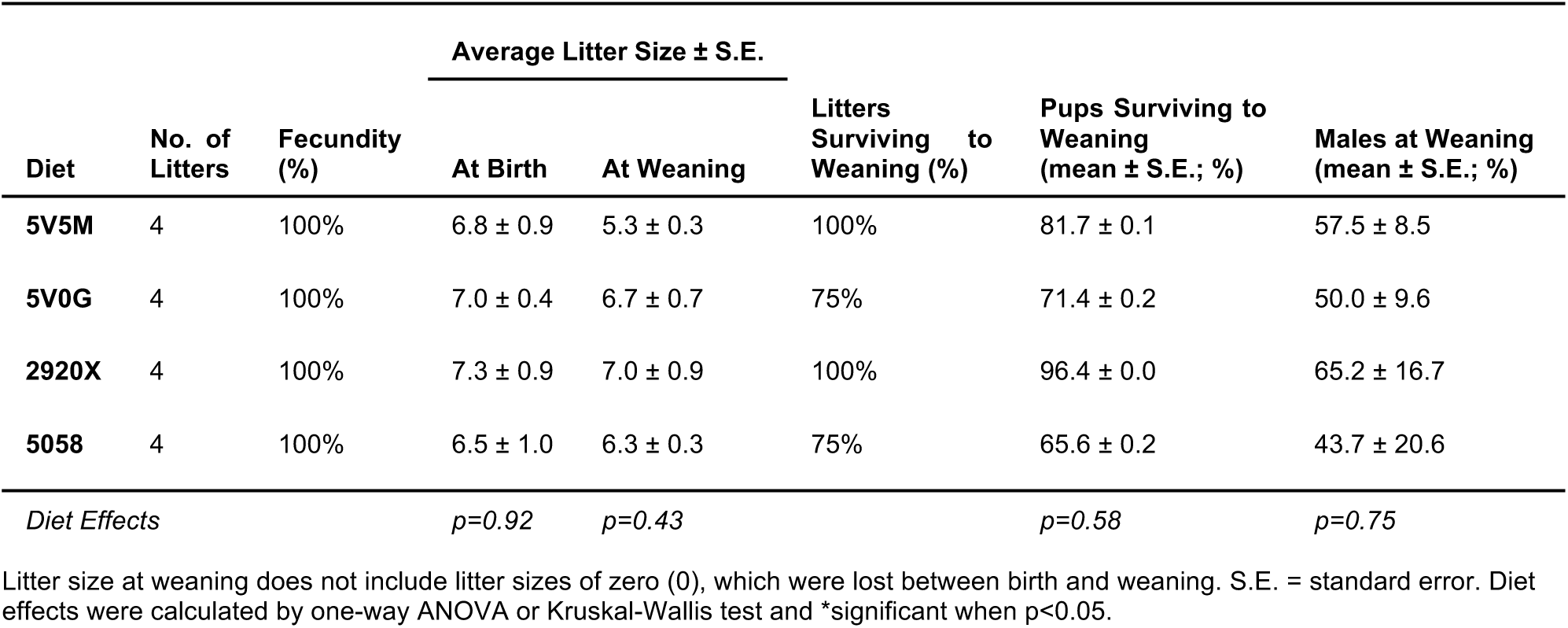
No effect of standard chow diets on AE model breeding success.

During development, when LE and DE mice were siblings housed together, there was a significant effect of chow diet on male and female body weight as early as postnatal day 5 (PND 5), where mice on the 5V5M diet showed 1.2× lower body weight compared to the other three diets (**Figure 5a**). By weaning, 5V5M-fed mice still weighed the least but were similar to 2920X, while 5V0G and 5058 exhibited higher body weights (**Figure 5b**). By adulthood, in the LE model (where mice remained on the different chows postweaning), 5V5M-fed mice still weighed the least, but a significant diet effect was only detected for male mice and was characterized by higher weights for 5058-fed mice (**Figure 5c**). In contrast, adult mice in the DE model exhibited no significant differences in body weight (**Figure 5d**). Crown-rump length (CRL), a measure of growth, followed similar (but not significant) body weight differences in the LE and DE model, except for adult LE females, where 5058-fed mice exhibited significantly shorter CRL (**Figure 5c**). Thus, exposure to different standard chow diets induces variability in mouse model developmental growth phenotypes in a sex-dependent manner but placing mice on the same diet at weaning reduces this variability.

**Figure 5.**
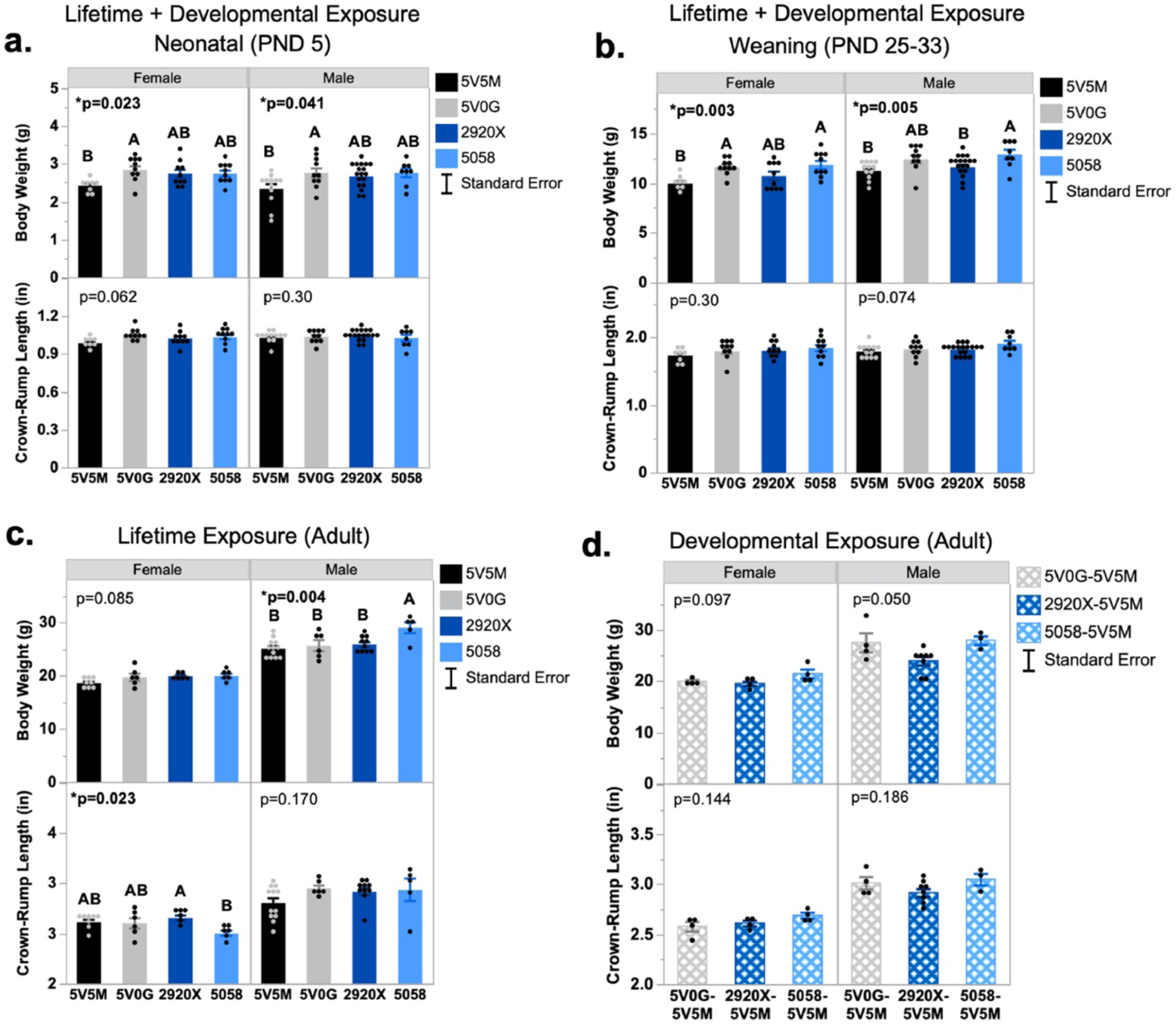
**Standard chow diets alter body weight and crown-rump length (CRL) in LE model but not DE model**. Body weight and CRL measures were measured on live offspring at **a.** neonatal (PND 5), **b.** weaning (PND 25-33), and **c-d**. adult (12 weeks) time points. LE and DE models were analyzed separately. Error bars represent the standard error of the mean for all graphs. Diet effects were calculated by one-way ANOVA or Kruskal-Wallis test and *significant when p<0.05.

To determine whether diet-induced differences in developmental growth phenotypes are associated with differences in adult phenotypes that are known metabolic health indicators, we measured effects on adult body composition, fat depot size, and glycemic status. In the LE model, 5058-fed females had 1.6× higher total fat mass than 2920X-fed females but no difference in lean mass (**Figure 6a**). In contrast, 5058-fed males had no difference in fat mass but significantly higher total lean mass than all other diets (**Figure 6a**), which mirrored their body weight differences (**Figure 5c**). In the DE model, we detected no significant effects of preweaning diet on fat mass in females or males (**Figure 6b**). However, consistent with the LE model, 5058-fed males in the DE model exhibited significantly higher total lean mass (**Figure 6b**) that mirrored the body weights (**Figure 5d**).

**Figure 6.**
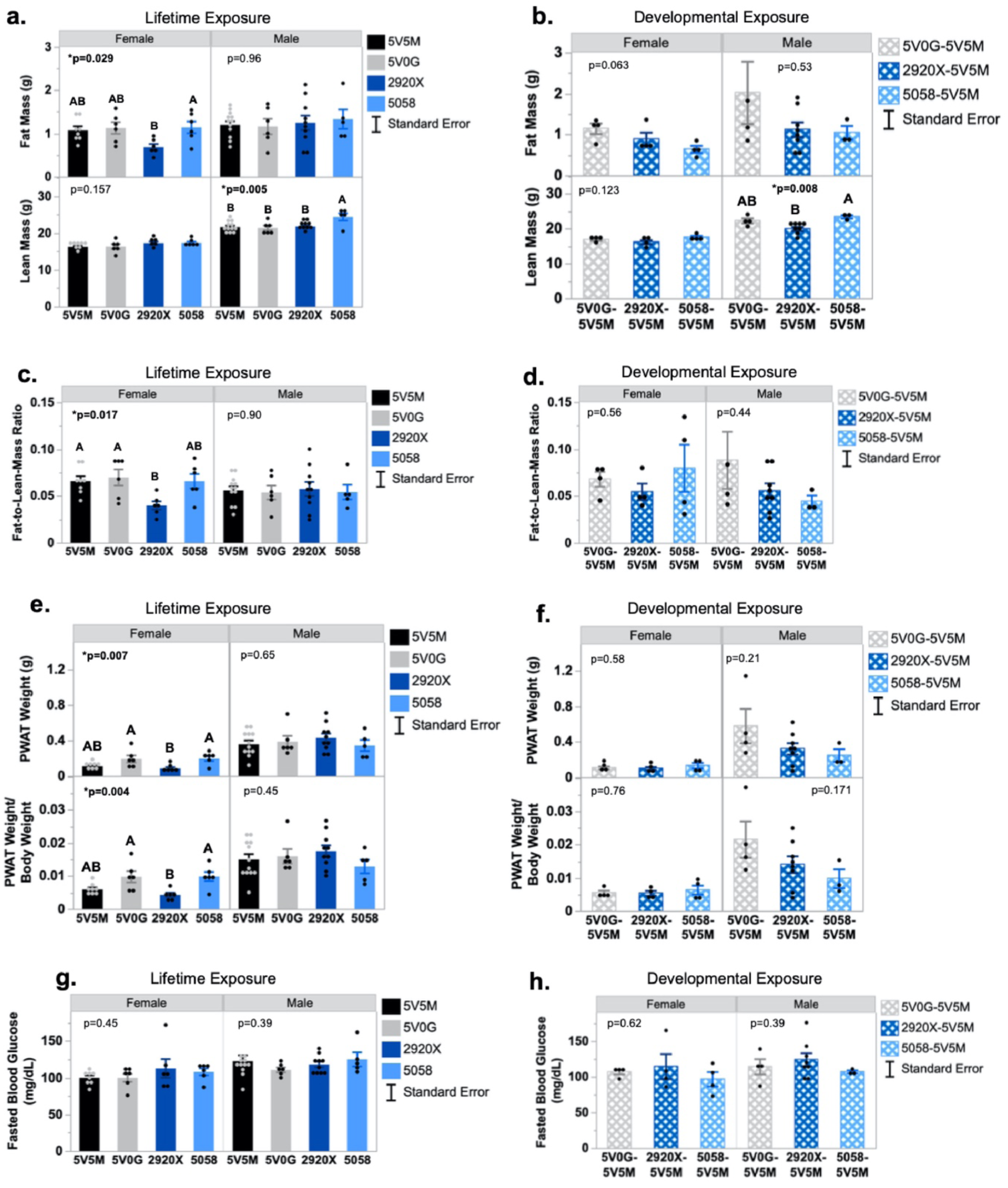
Standard chow diets alter adult body composition and metabolic health outcomes for LE and DE models. **a-d.** EchoMRI measured body composition for 8-9-week-old female and male offspring. Total fat and total lean mass were measured for **a.** LE and **b.** DE models. Fat-to-lean-mass ratios were calculated for **c.** LE and **d.** DE models. **e-h.** Perigonadal fat pad weights (PWAT) were collected for **e.** LE males (12-14 weeks) and females (15-16 weeks) and **f.** DE males (12-14 weeks) and females (15-16 weeks). Fasted whole blood glucose was measured in live **g.** LE males (12-14 weeks) and females (15-16 weeks) and live **h.** DE males (12-14 weeks) and females (15-16 weeks). LE and DE models were analyzed separately. Error bars represent the standard error of the mean for all graphs. Diet effects were calculated by one-way ANOVA or Kruskal-Wallis test and *significant when p<0.05.

Total fat-to-lean-mass ratios are considered a predictor of metabolic disease, and lower ratios are considered protective. In comparison, higher ratios suggest an increased risk of cardiac events and death^19^. In the LE model, diet drove significant differences in fat-to-lean-mass ratios in females, with 2920X-fed mice exhibiting ∼1.7× lower ratios than mice fed the other diets (**Figure 6c**). The DE model did not exhibit any diet-induced differences in fat-to-lean-mass ratios, although 2920X-fed females had the lowest ratios (**Figure 6d**).

The perigonadal fat pad is usually the largest white adipose tissue depot in the mouse, and larger size is associated with impaired metabolic health^20^. In the LE model, 5058-and 5V0G-fed females exhibited ∼2× larger PWAT weights than 2920X-fed mice, which remained significant even after normalizing to body weight (**Figure 6e**). These effects partly mirrored total body fat mass differences (**Figure 6a**). There were no significant diet-induced differences in PWAT measurements for LE males (**Figure 6e**) or DE males or females (**Figure 6f**). There were also no diet effects on glycemic status in the LE model (**Figure 6g**) or DE model (**Figure 6h**).

Consistent with growth phenotypes, these findings show chow diet differences can drive variability in metabolic health phenotypes that is sex-specific and may be ameliorated by weaning mice onto the same diet.

### Mice with lifetime exposure to different standard chows are more susceptible to altered composition of the gut bacterial microbiota

To investigate the impact of different standard chow diets on the composition of the gut bacterial microbiota, we performed 16S rRNA sequencing of DNA from cecal contents. Since mice in the LE model exhibited the greatest phenotypic response, we assessed microbiota changes in this group compared to the parental AE model, which exhibited no detectable phenotypic response.

Alpha diversity describes microbial richness and how evenly distributed microbes are within a sample, and the Shannon index is used as a measure of alpha diversity^21^. In the AE model, females exhibited significant diet-induced differences in microbial richness, with 5058-fed females exhibiting the highest number of observed sequence variants and 2920X-fed mice exhibiting the lowest (**Figure 7a**). AE males were not similarly affected (**Figure 7a**), and there were no diet effects on the AE model’s Shannon index (**Figure 7b**). In contrast, LE mice exhibited no diet effects on microbial richness (**Figure 7c**), but diet significantly altered the Shannon index, with 5058-and 5V0G-fed mice exhibiting significantly lower Shannon indices than 2920X-fed mice (**Figure 7d**).

**Figure 7.**
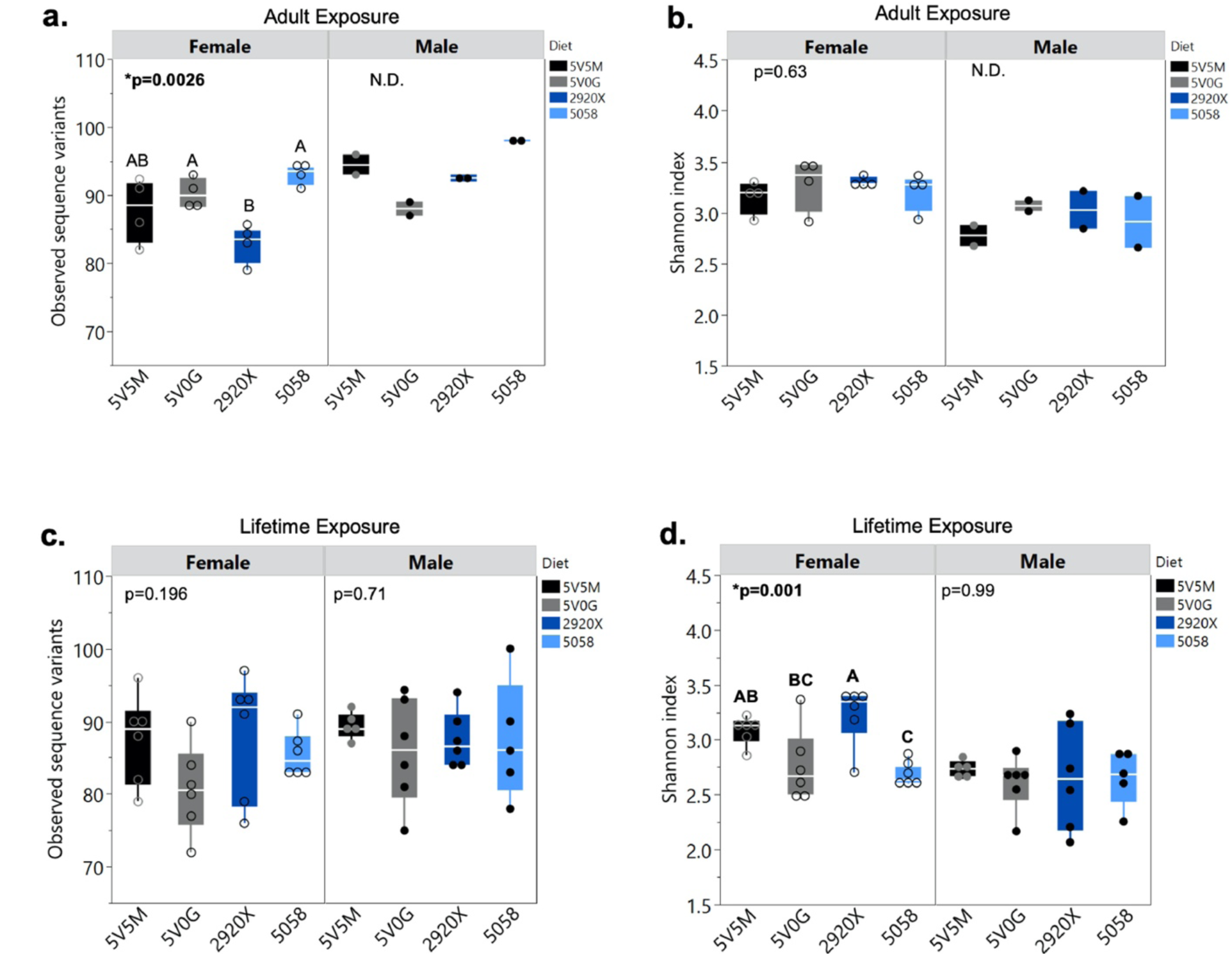
Standard chow diet alters the alpha diversity of gut bacterial microbiota for AE and LE models. AE Model: **a.** Observed sequence variants and **b.** Shannon index. LE Model: **c.** Observed sequence variants and. **d.** Shannon index. Box plot elements include: median (center line), upper 75^th^ quartile and lower 25^th^ quartile, and expected variation of the data (whiskers). Open circles indicate female samples, and closed circles indicate male samples. Main diet effects were calculated by linear regression and *significant when p<0.05. N.D. = p-value not determined due to small sample size.

Beta diversity describes the variability in microbes across samples within a population^21^. The effects of chow diets on the beta diversity of the gut bacterial microbiota were determined using principal coordinate analysis (PCoA) and PERMANOVA with Bray-Curtis distances. In the AE model, diet and sex drove significant differences in beta diversity, with PC1 explaining 25.1% of the variability and PC2 explaining 13.1% (**Figure 8a**). Among the four diets, 5058-fed mice clustered furthest away from the other three (**Figure 8a**). There was no *sex × diet* interaction effect, and when the samples were stratified by sex, there was a similar clustering of diets but no detectable diet effect, likely due to the small sample size (**Figures 8b-c**). In the LE model, diet drove significant differences in beta diversity that were similar to the AE model, with similar amounts of variance explained on PC1 and PC2 (23.2% and 11.6%, respectively) and with distinct clustering of 5058-fed mice (**Figure 8d**). However, there was also a significant *sex* and *sex × diet* interaction effect detected in the LE model (**Figure 8d**). When stratified by sex, females and males exhibited a significant diet effect, with 5058-fed mice remaining distinct from mice fed the other diets (**Figure 8e-f**).

**Figure 8.**
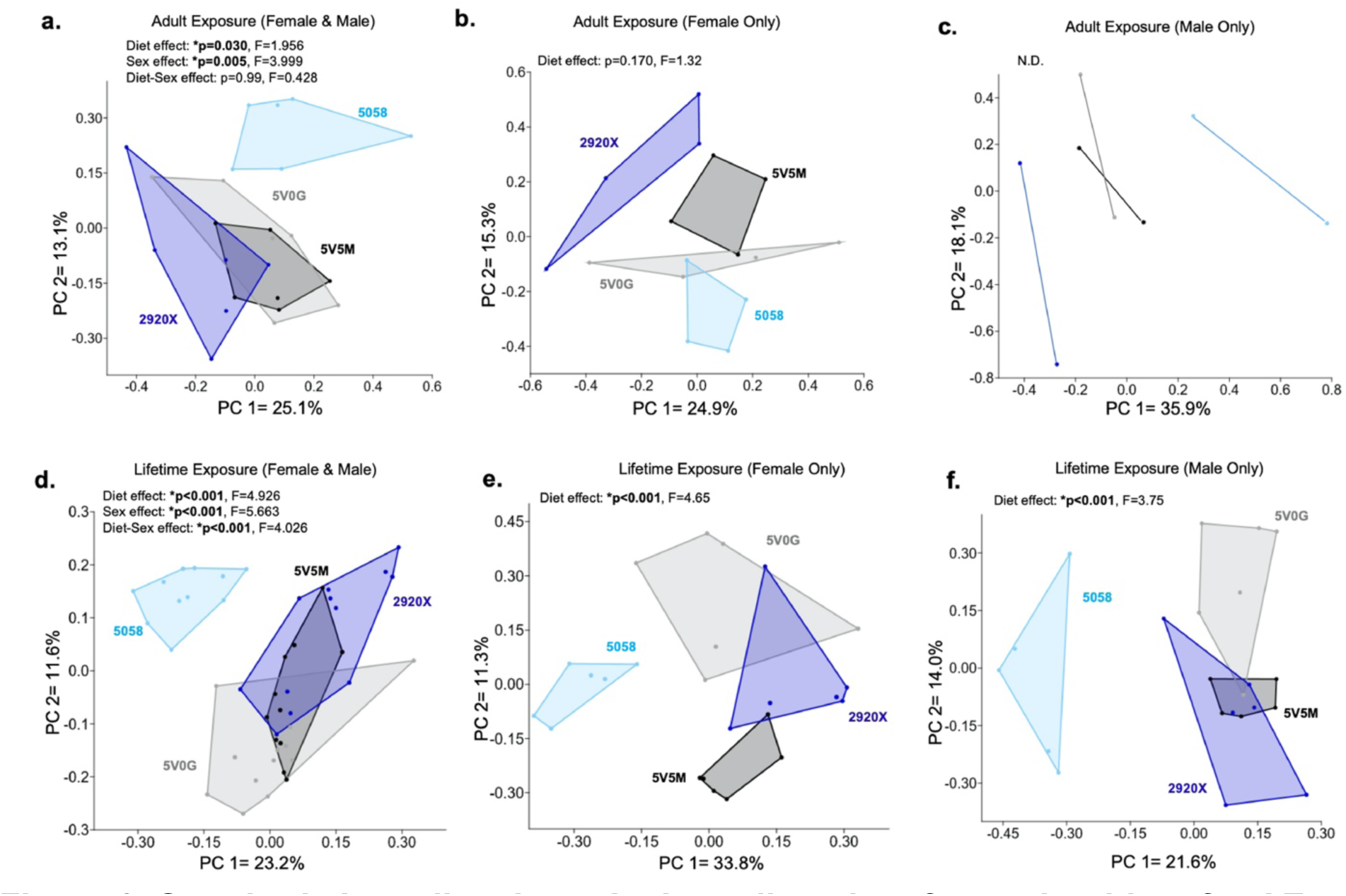
Standard chow diet alters the beta diversity of gut microbiota for AE and LE models. AE Model: **a.** PCoA using two-way PERMANOVA and Bray-Curtis distances and **b-c.** PCoA using one-way PERMANOVA and Bray-Curtis distances. LE Model: **d.** PCoA using two-way PERMANOVA and Bray-Curtis distances and **e-f.** PCoA using one-way PERMANOVA and Bray-Curtis distance. Main diet or sex effects were calculated by linear regression and *significant when p<0.05. N.D. = p-value not determined due to small sample size.

Finally, we investigated the impact of diet on the differential relative abundance of individual bacterial genera using a multi-factor model (diet + sex) and validated using ANCOM-BC2^22^ to identify diet effects after adjustment for sex effects. In the AE model, chow diet significantly altered the abundance of three bacterial genera (*Clostridium sensu stricto 1*, *Ruminococcus*, and *Tuzzerella*) similarly for males and females (**Figure 9a-c**). ANCOM-BC2 confirmed these effects. Genera altered by AE to the diets were also differentially abundant in the LE model but with differences between the sexes (**Figure 9d-f**). 5058-fed mice exhibited an increased abundance of *Clostridium sensu stricto 1* in AE males and females, but only females were affected in the LE model (**Figure 9a** & **d**). 5058-and 5V0G-fed mice also exhibited an increased abundance of *Ruminococcus* in AE males and females, but only males were affected in the LE model (**Figure 9b** & **e**).

**Figure 9.**
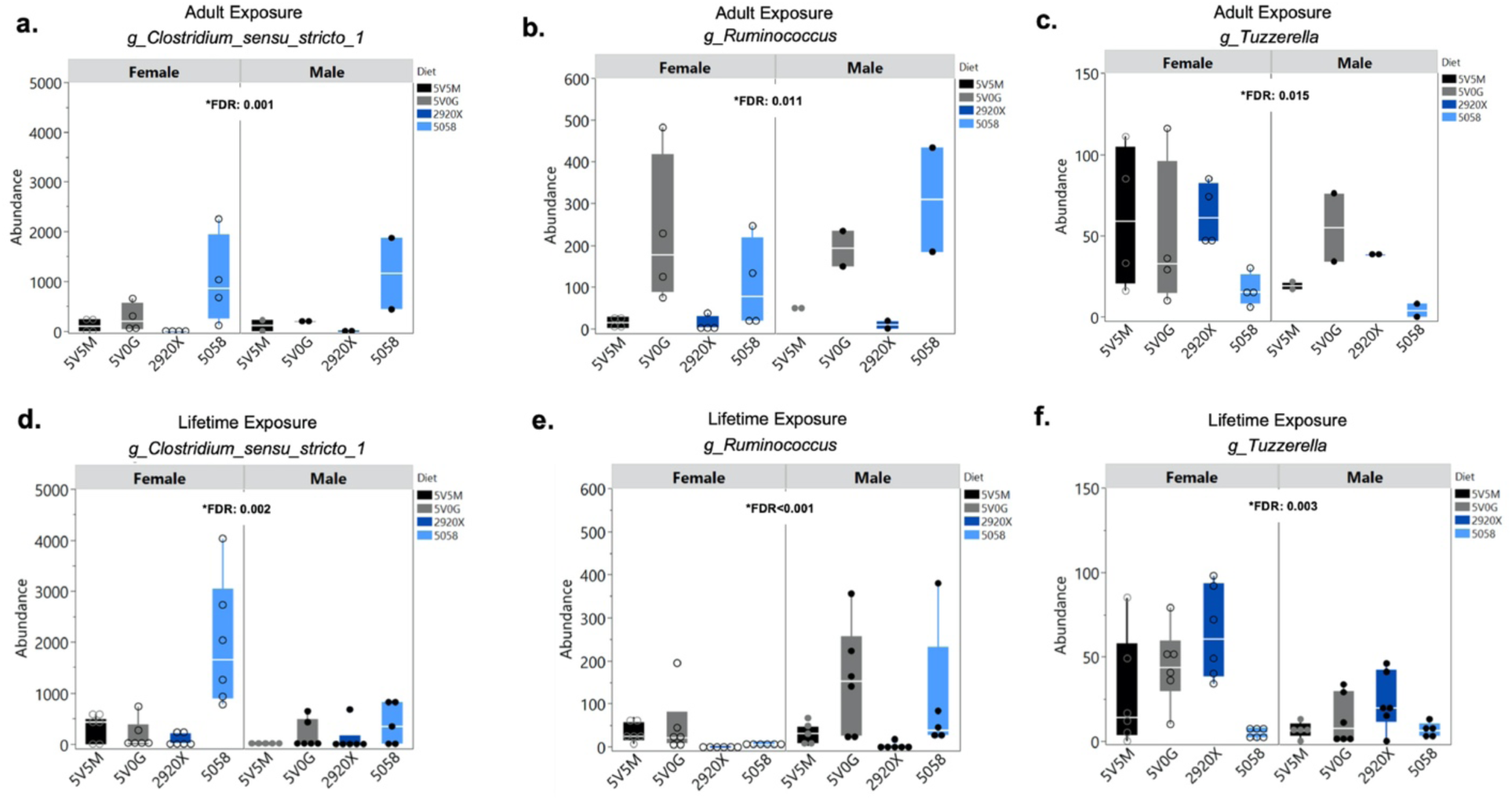
Differentially abundant bacterial genera affected in both AE and LE models after adjusting by sex. **a-c.** Diet-dependent abundance of genus (*Clostridium_sensu_stricto_1*, *Ruminococcus*, & *Tuzzerella*) for AE model and **d-f.** LE model after multi-factor analysis of microbial differential abundance with “diet” as the primary effect and “sex” as a covariate. Box plot elements include: median (center line), upper 75^th^ quartile and lower 25^th^ quartile, and expected variation of the data (whiskers). Open circles indicate female samples and closed circles indicate male samples. FDR (False Discovery Rate) shown for main diet effects after adjustment for sex and *significant when FDR<0.05.

On the other hand, 5058-fed mice exhibited the lowest abundance of *Tuzzerella* in males and females of both models, but overall abundance was substantially lower for all diets in the LE model versus the AE model (**Figure 9c** & **f**). ANCOM-BC2 confirmed the differential abundance of all genera except *Ruminococcus* (**Figure 9b** & **e**), which yielded *p*<0.05, but failed to pass bias correction.

The LE model exhibited significant diet effects on the abundance of 24 additional bacterial genera (27 total, **Table 2**). ANCOM-BC2 confirmed all except three genera (*Muribaculaceae*, *Eubacterium xylanophilum* group, and *Eubacterium ventriosum* group) which failed to pass correction for multiple testing. Interestingly, females showed larger differences in genus abundance with 5058-and 2920X-fed females exhibiting the greatest effect sizes (**Table 2**). Two-way hierarchical clustering of all 27 genera confirmed that samples clustered in part by diet, with the most distinct clusters observed for 5058-and 2920X-fed females in the LE model (**Supplemental Figure 6**). This distinct clustering is seemingly driven by five genera with greater abundance in 5058-fed mice and lower abundance in 2920X-fed mice (*Eubacterium xylanophilum* group, *Clostridium sensu stricto 1*, *Lactobacillus*, *Bifidobacterium*, and *Eubacterium nodatum* group) (**Table 2**, **Supplemental Figure 6**); and 12 bacterial genera with lower abundance in 5058-fed mice but higher abundance in 2920X-fed mice (*Anaeroplasma*, *Tuzzerella, Eubacterium ventriosum* group*, Acetatifactor*, *Blautia, Clostridia* vadinBB60 group, *Alistipes*, *Colidextribacter*, *Oscillibacter*, *Intestinimonas*, *Lachnospiraceae* FCS020 group, and *Butyricicoccaceae* UCG-009) (**Table 2**, **Supplemental Figure 6**).

**Table 2.**
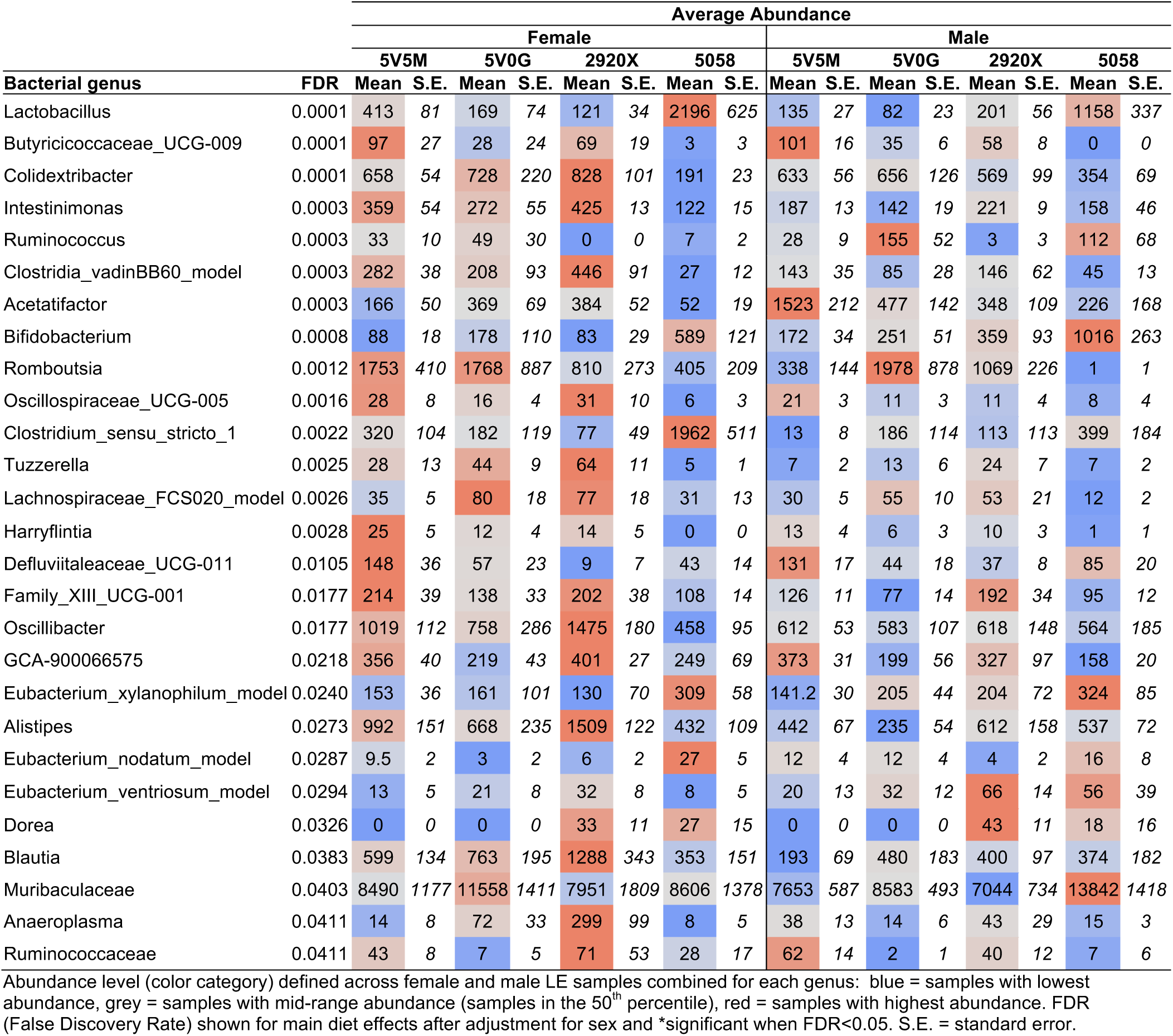
Standard chow diet alters the differential abundance of microbes in LE model.

## Discussion

In this study, we demonstrate that the timing of exposure to different chow diets determines the extent of diet impact on mouse model phenotypes even among genetically identical animals. While a few previous studies have measured the phenotypic impact of adult exposure to standard chow^9,11,23^, this is the first study to quantitatively measure the differences in nutrient intake among chow diets and test the impact of exposure during different life stages, including adult (AE), lifetime (LE), and developmental exposure (DE). Furthermore, including both sexes in our study provides new evidence of sex-dependent diet effects, which have been poorly assessed in previous literature. Ultimately, this study highlights the important roles of the chow diet, timing of exposure, and sex in the reproducibility of preclinical mouse model phenotypes. Our finding that feeding mice a common diet postweaning ameliorated many phenotypes implicates this as a viable method for improving phenotypic reproducibility across studies where mice may need to be bred on different chow diets. Chow diet effects on the gut bacterial microbiota were also investigated. Surprisingly, despite dissimilar phenotypic responses, the AE model exhibited a significant but limited microbial response. LE mice exhibited a broader microbial response that encompassed effects observed in the AE model.

Mice fed different standard chows exhibited different food intake levels, and these differences substantially impacted nutrient exposure. The largest effects were observed for 5058 mice, with the highest adult food intake resulting in higher macronutrient and fiber exposure levels. These higher intake levels likely contributed to the larger body weights observed in 5058-fed mice in the LE model.

While most chow diet studies highlight the effects of calorie and macronutrient differences, our study adds new information about vitamin D and folic acid content, two micronutrients that are widely variable among chow diets. During development, vitamin D supports a healthy *in utero* environment and regulates offspring muscle mass, bone health, adipose storage, and brain development^24^. In this study, the vitamin D content of all four diets exceeded the recommended levels, which are already documented as potentially above the required amount^16^. Folate plays critical roles in DNA synthesis and 1-carbon metabolism but is most well-recognized for regulating offspring neural tube formation^25^. In this study, folic acid content in the 5V0G diet was >4× higher than the recommended amount^17^ putting it close to experimentally supplemented diets^18^. Despite the high variability in micronutrient intake across the diets for all exposure models, only the LE model exhibited a significant difference in circulating levels of vitamin D (25(OH)D). Circulating folate concentrations were high but similar among the models. The fact that the LE model had the longest exposure period (3 weeks prenatal, 3 weeks preweaning, and 9-12 weeks postweaning) implicates the length of chow diet exposure in driving differences in micronutrient status.

Timing of exposure also determined phenotypic response. Mice in the LE model demonstrated heightened sensitivity to nutrient variability compared to AE and DE models. For example, 5V5M-fed mice exhibited restricted body weight across the lifespan in the LE model. On the other hand, 5058-fed males in the LE model revealed connections between chow diet-driven increased food intake, nutrient intake, adult body weight, and adult lean mass. 2920X-fed females in the LE model exhibited restricted PWAT, growth, and reduced total body fat & fat-to-lean mass ratios. These effects were not observed in the AE or DE models. Thus, diet timing influences the extent of impact on phenotype reproducibility. Chow diet selection and use are critical in studies on developmental growth phenotypes and body composition phenotypes commonly used as indicators of metabolic health in mice. Importantly, a normalization diet postweaning may be an effective way to reduce unwanted diet-induced phenotypic variability in mouse models.

This is the first study to demonstrate that the timing of chow diet exposure and sex influence variability in the gut-bacterial microbiota composition. Nutrients are well known to play a major role in the establishment and maintenance of the gut microbiome^26^. Here, we showed that differences in chow diet drive changes in the gut’s bacterial composition regardless of whether exposure was during development + adulthood (LE) or only in adulthood (AE). However, while the diet effects on beta diversity and differential abundance of bacterial genera in the AE model were mostly replicated in the LE model, the LE model exhibited 9× more differentially abundant genera that were sex-dependent. Among the diets, 5058 exhibited the most distinct effect on beta diversity, while both 5058 and 2920X stood out as the extremes in diet effects on bacterial genera abundance. Although we were unable to measure the phenotypic consequences of these microbial differences, several have predicted pathogenic effects (*Clostridium sensu stricto 1*^27^*, Ruminococcus*^28^*, Tuzzerella*^29^, *Acetatifactor*^30^) while others are predicted to have protective effects on a range of outcomes including gut and metabolic health (*Bifidobacterium*^31^*, Lactobacillus*^32^*, Eubacterium nodatum* group, *Alistipes*^33^*, Blautia*^34^*, Eubacterium xylanophilum* group^35^).

This study has methodically tested several new parameters of chow diet exposures and their effects on experimental reproducibility. Our findings highlight the importance of standard chow selection and the timing of its use in experimental mouse model research. This further underscores the necessity of reporting the exact diets used and the timing and duration of dietary exposures within preclinical mouse model studies. Thus, rigorous study design and reproducibility of outcomes require the development of specific guidelines on required reporting of diet expanded beyond what has already been initiated *via* valuable resources like the ARRIVE Guidelines^36^.

Here, we focused on macro-and micronutrient differences among the diets, while recognizing that other dietary components, such as bioactive compounds, could also be influencing the observed outcomes. Importantly, although diet effects were strikingly similar between males and females for measures taken before weaning, all significant timing and diet-dependent adult/postweaning phenotypes were sex-specific. Despite the study only offering a limited assessment of males in the AE model (due to low sample sizes), the extensive data provided for the other models further underscores the need to include both sexes in studies of factors affecting experimental reproducibility.

## Methods

### Animal husbandry, dietary treatment, and breeding

All animals were handled in accordance with the Guide for the Care and Use of Laboratory Animals under an approved and registered animal use protocol at the University of North Carolina (UNC) at Chapel Hill. Throughout the study, all mice were housed in ventilated microisolator cages with corncob bedding and maintained at a vivarium temperature of 21-23°C with a 12-hr light cycle and *ad libitum* access to sterilized water and standard chow. Animals were monitored daily for stress and adverse health. Any animals that demonstrated stress, adverse health, and/or did not survive until euthanasia were excluded from the study following established animal care guidelines.

#### Standard chow selection

Four standard chows were selected for this study: 5V5M (PicoLab Select Mouse 50 IF/9F; Lab Diet, Durham, NC), 5V0G (Select Mouse 50 IF/9F Auto; Lab Diet, Durham, NC), 2920X (Teklad Global Soy Protein-Free Extruded Rodent Diet; Teklad, Madison, WI), and 5058 (PicoLab Mouse Diet 20; Lab Diet, Durham, NC). Diets were selected based on the high prevalence of use at UNC and >18 donor institutions that submit mice to the UNC Mutant Mouse Resource & Research Center strain repository^37^.

#### Adult exposure model (AE)

Sample sizes are provided in **Supplemental Table 3**. Six-week-old virgin female and male C57BL/6J mice were simultaneously procured from Jackson Laboratories (stock# 000664, Bar Harbor, ME) and transferred with random assignment (separated by sex) into cages with one of four standard chow diet groups (5V5M, 5V0G, 2920X, & 5058) (**Figure 1f**). After 3 weeks of acclimation to dietary treatment, a subset of females was euthanized to collect pre-mating micronutrient measurements (**Figure 1f**). The remaining mice were mated in trio breeding for 3 weeks. Bred males (sires) were euthanized after breeding and bred females (dams) were euthanized at weaning (**Figure 1f**). Time on a diet is provided in **Supplemental Table 4**. All mice were fasted 12-hrs before euthanasia by CO_2_.

#### Lifetime exposure (LE) and Developmental exposure (DE) models

Sample sizes are provided in **Supplemental Table 5**. At weaning, pups were caged by sex, and siblings were assigned to either remain on their preweaning diet for the lifetime exposure (LE) or switch to a common postweaning diet (5V5M) for the developmental exposure (DE) (**Figure 1f**). At the end of treatment, all mice were fasted for 12-hrs before euthanasia by CO_2_. Time on a diet is provided in **Supplemental Table 4**.

### Nutrient intake

Food intake was measured weekly for each individual cage, and average daily values were calculated per mouse [Avg ((food weight at week end - food weight at week start)/(# of days)/(# of mice in the cage))]. Each diet was vacuum-sealed separately and stored at -80°C. Diets were shipped overnight on dry ice to be measured by Eurofins Food Chemistry Testing (Madison, WI) (**Supplemental Table 1**). “*Measured values*” of total calories, macronutrients (fat, carbohydrates, & protein), fiber (total dietary fiber & crude fiber), and micronutrients (vitamin D2, vitamin D3, & folic acid) were converted into units of measures comparable to the “*Manufacturer’s estimates*”.

### Phenotype measurements

For the AE model, body weights were measured every 3 weeks after the start of treatment (pre-mating) and then once at the end of treatment. For the LE and DE models, body weights were measured biweekly, starting at postnatal day 5 (PND 5) until the end of treatment. Minimal handling was performed at sensitive periods (e.g., during gestation, birth-PND 4). Crown-rump length (CRL) was measured using carbon fiber calipers (Cat#36934-152, VWR) at PND 5, weaning (PND 25-33), and adulthood (PND 84). Body composition was measured at 8-9-weeks of age by EchoMRI (Animal Metabolism Phenotyping Core, University of North Carolina at Chapel Hill, Chapel Hill, NC). Fasting blood glucose was measured at the end of treatment after a 12-hr fast using a single tail nick bleed to measure blood glucose levels *via* glucometer (Cat# 68623221868, Accu-Chek Performa, Roche). Perigonadal white adipose tissue (PWAT) was dissected and weighed at the end of treatment immediately following euthanasia.

### Serum measurements (vitamin D & folate)

Serum was isolated from whole blood collected by cardiac puncture and centrifuged 10 min × 2,000 g at 4°C. Serum was flash-frozen in liquid nitrogen and stored at −80°C until use. Vitamin D (25(OH)D) levels were measured by ELISA following the manufacturer’s protocol (Mouse/Rat 25-OH Vitamin D ELISA Assay Kit (VID21-K01), Eagle Biosciences, Nashua, NH). Serum folate concentrations were determined using the *Lactobacillus casei* microbiological assay as previously described^38^.

### Statistical analyses for phenotype data

Unless otherwise stated, statistical analyses were performed using JMP Pro 17 Software (SAS, NC). Data were tested for normality (Shapiro-Wilk test), and normally distributed data were analyzed using one-way ANOVA. Non-normally distributed data were analyzed using the Kruskal-Wallis test. Tukey’s post-hoc was used to identify which diets differed significantly.

### Bacterial microbiota profiling

Cecum was harvested after euthanasia from AE females and males (n = 4/diet & n = 2/diet) and LE females & males (n = 6/diet & n = 5-6/diet), flash-frozen in liquid nitrogen, and stored at -80°C. Samples close to the population median for body weight, body composition, and cecum weight in each diet group were selected from 3-4 litters/diet.

Frozen samples were shipped overnight on dry ice to the University of Missouri (MU) Metagenomics Center (Columbia, MO) for DNA extraction, prep, and plating. 16S rRNA library preparation and sequencing were performed at the MU Genomics Technology Core, data cleanup and QC, sequence alignments, and annotations at the MU Bioinformatics and Analytics Core, as previously described^11^. In brief, bacterial 16S rRNA amplicons were built via amplification of the V4 region of the 16S rRNA gene using the U515F/806R primer pair^39^ and processed for paired-end sequencing by MiSeq (Illumina) as previously described^11,39,40^. Read pairs were rejected if either read did not match the 5’ primer (using an error rate of 0.1). Cutadapt^41^ (version 2.6) was used to trim primers. Obtained sequence data were de-multiplexed using the QIIME2^42,43^ DADA2^43^ plugin (version 1.10.0) as follows: forward and reverse reads were truncated to 150 bases; forward and reverse reads with expected errors >2.0 were discarded; and chimeras were detected using the "consensus" method and removed. Output sequences were classified as amplicon sequence variants (ASVs). Taxonomy was assigned using the SILVA.v132^44^ reference database with the classify-sklearn procedure. The ASV dataset (**Supplemental Excel Table 1**) was randomly subsampled to a uniform sequence depth of 42,577 reads per sample for rarefaction. The rarefied table (**Supplemental Excel Table 2)** contained 2,980,390 reads (42,577/sample) and 205 OTUs.

Alpha and beta diversity metrics were calculated in PAST4.13^45^ using rarefied data. Linear regression models and Tukey’s post-hoc tests were used to measure main diet effects on observed sequence variants and Shannon index using JMP Pro 17 Software (SAS, NC). Permutational multivariate analysis of variance (PERMANOVA) using Bray-Curtis distances was performed to test for “Diet” and “Sex” effects, and data were quarter-root transformed to generate the Principal Coordinate Analysis (PCoA). Microbial differential abundance was calculated in MicrobiomeAnalyst 2.0 (available at “microbiomeanalyst.ca”) using the Marker Data Profiling tool. Features with identical values (i.e. zero) across all samples and features appearing in only one sample were excluded, resulting in a total of 2,980,202 reads (average of 42,574 reads/sample) and 205 ASVs analyzed. Multi-factor comparisons with the linear model at the genus level were performed for female and male samples combined to evaluate “Diet” as the primary effect and “Sex” as a covariate. Filtering was applied to features with low count (<4), low prevalence (<20%), and low variance (<10% based on inter-quantile range). FDR<0.05 after Bonferroni^46^ and Benjamini-Hochberg^47^ corrections were considered statistically significant. Confirmatory differential abundance analysis was performed using Analysis of Compositions of Microbiomes with Bias Correction 2 (ANCOM-BC2). ANCOM-BC2 testing was performed using fixed effects for "Diet" and "Sex”. Taxa exhibiting significant differential abundance were determined by a Benjamini-Hochberg^47^ corrected p-value of less than 0.05 (**Supplemental Excel Tables 3-4**). Structural zeroes were identified based on the presence or absence of taxa across the groups defined by the “Diet” variable.

Two-way hierarchical clustering of differentially abundant microbes at the genus level was performed in JMP Pro 17 Software (SAS, NC). AE and LE groups were analyzed separately.

## List of Abbreviations

AE: adult exposure
LE: lifetime exposure
DE: developmental exposure

## Acknowledgments

We thank Michael Whalen from the Ideraabdullah Lab for input on the paper format, Jackie Brooks for providing chow diet stats from the UNC MMRRC, and Fernando Matias from the MacFarlane Lab for technical support.

## Author Contributions

**Megan M. Knuth** – Writing – Original Draft, Methodology, Formal Analysis, Investigation, Data Curation, Visualization**; Carolina Vieira Campos** – Writing – Original Draft, Formal Analysis, Data Curation, Visualization; **Kirsten Smith** – Formal Analysis, Investigation, Data Curation, Writing – Editing & Review; **Elizabeth K. Hutchins** – Investigation, Data Curation, Writing – Editing & Review; **Shantae Lewis** – Data Curation, Writing – Editing & Review; **Mary York** – Formal Analysis; **Lyndon M. Coghill** – Formal Analysis; **Craig Franklin** – Advice on Methodology & Formal analyses, Writing – Editing & Review; **Amanda MacFarlane** – Methodology, Investigation, Writing – Editing & Review; **Aaron Ericsson** – Methodology, Formal Analysis, Software, Investigation, Data Curation, Writing – Editing & Review; **Terry Magnuson** – Conceptualization, Methodology, Resources, Supervision, Funding Acquisition, Writing – Editing & Review; **Folami Ideraabdullah** – Writing – Original Draft, Conceptualization, Methodology, Investigation, Resources, Data Curation, Supervision, Project Administration, Funding Acquisition.

## Data Availability

Data will be made fully available upon request. Microbiome OTU datasets (unrarefied & rarefied) and ANCOM-BC2 data are provided in **Supplemental Excel Tables 1-4**. Additional data will be made available upon request.

