## Supplemental Data_Tables & Figures for "Timing of standard chow exposure determines the variability of mouse phenotypic outcomes and gut microbiota profile"

**Affiliations**:
[1]Department of Genetics, School of Medicine, University of North Carolina at Chapel Hill, Chapel Hill, NC 27599, USA;
[2]Lineberger Comprehensive Cancer Center, University of North Carolina at Chapel Hill, Chapel Hill, NC 27599, USA;
[3]Department of Translational Medicine, School of Medical Sciences, State University of Campinas, Campinas 13083-881, Brazil;
[4]Postbaccalaureate Research Education Program, University of North Carolina at Chapel Hill, Chapel Hill, NC 27599, USA;
[5]University of Missouri (MU) Bioinformatics and Analytics Core, Bond Life Sciences Center, Columbia, MO 65201, USA;
[6]Institute for Data Science and Informatics, University of Missouri, Columbia, MO 65211, USA;
[7]Department of Veterinary Pathobiology, MU, Columbia, MO 65201, USA;
[8]Mutant Mouse Resource and Research Center at the University of Missouri (MU MMRRC), Columbia, MO 65201, USA;
[9]MU Metagenomics Center (MUMC), University of Missouri, Columbia, MO 65201, USA;
[10]Texas A&M Agriculture, Food, and Nutrition Evidence Center, Fort Worth, TX 76102, USA;
[11]Department of Nutrition, Texas A&M University, College Station TX 77843, USA;
[12]Mutant Mouse Resource and Research Center, University of North Carolina at Chapel Hill, Chapel Hill, NC 27599, USA;
[13]Department of Nutrition, Gillings School of Global Public Health, University of North Carolina at Chapel Hill, Chapel Hill, NC 27599, USA;

***Corresponding Author**:
Folami Ideraabdullah


**List of Abbreviations**: **AE** – adult exposure; **LE** – lifetime exposure; **DE** – developmental exposure.

### Tables

#### Supplemental Table 1. Measured nutrient values for key macro- & micronutrients (Eurofins).

| Diet | Calories from Carbs (%) | Calories form Fat (%) | Calories from Protein (%) | Total Calories (Kcal/g) | Crude Fiber (%) | Folic Acid (mg/kg) | Vitamin D3 (IU/g) | Vitamin D2 (IU/g) |
|---|---|---|---|---|---|---|---|---|
| **5V5M** | 59.60 | 25.39 | 19.43 | 3.81 | 2.14 | 2.01 | 1.79 | 1.92 |
| **5V0G** | 61.54 | 24.66 | 19.10 | 3.71 | 2.87 | 8.09 | 3.13 | 1.07 |
| **2920X** | 69.29 | 15.83 | 21.10 | 3.53 | 2.59 | 1.96 | 1.23 | 1.14 |
| **5058** | 58.21 | 24.70 | 22.24 | 3.67 | 2.44 | 2.34 | 2.73 | 0.67 |

#### Supplemental Table 2. Manufacturer's estimates for key macro- & micronutrients (Lab Diet & Teklad).

| Diet | Calories from Carbs (%) | Calories form Fat (%) | Calories from Protein (%) | Total Calories (Kcal/g) | Crude Fiber (%) | Folic Acid (mg/kg) | Vitamin D3 (IU/g) |
|---|---|---|---|---|---|---|---|
| **5V5M** | 58.35 | 21.42 | 20.24 | 3.61 | 2.30 | 3.6 | 2.3 |
| **5V0G** | 58.01 | 21.54 | 20.45 | 3.58 | 2.30 | 8.6 | 2.3 |
| **2920X** | 60.00 | 16.00 | 24.00 | 3.10 | 2.70 | 4.0 | 1.5 |
| **5058** | 55.23 | 21.56 | 23.21 | 3.45 | 2.40 | 2.9 | 3.4 |

### Supplemental Table 3. Sample sizes for AE model.

| | | | | | | Diet Acclimation | | | Breeding | | |
|---|---|---|---|---|---|---|---|---|---|---|---|
| Model | Sex | Diet | Total # of Cages | Total # of Mice in Each Cage | Total # of Mice | # of Cages Euthanized at Mating | # of Mice in Each Cage Euthanized at Mating | # of Mice Euthanized at Mating | Total # of Breeding Cages | Total # of Mice Mated in Each Breeding Cage | Total # of Mice Bred |
| Adult Exposure | Female | 5V5M | 2 | 4, 5 | 9 | 1 | 5 | 5 | 2 | 2, 2 | 4 |
| | | 5V0G | 2 | 4, 5 | 9 | 1 | 5 | 5 | 2 | 2, 2 | 4 |
| | | 2920X | 2 | 4, 5 | 9 | 1 | 5 | 5 | 2 | 2, 2 | 4 |
| | | 5058 | 2 | 4, 5 | 9 | 1 | 5 | 5 | 2 | 2, 2 | 4 |
| | Male | 5V5M | 1 | 2 | 2 | 0 | 0 | 0 | 2 | 1, 1 | 2 |
| | | 5V0G | 1 | 2 | 2 | 0 | 0 | 0 | 2 | 1, 1 | 2 |
| | | 2920X | 1 | 2 | 2 | 0 | 0 | 0 | 2 | 1, 1 | 2 |
| | | 5058 | 1 | 2 | 2 | 0 | 0 | 0 | 2 | 1, 1 | 2 |

### Supplemental Table 4. Time on diet for AE, LE, and DE models.

| Model | Sex | Diet | Time on Diet Before Birth (Weeks) | Time on Diet from Birth until Weaning (Weeks) | Time on Diet Postweaning (Weeks) | Total Time on Diet (Weeks) |
|---|---|---|---|---|---|---|
| | | | Estimated | Mean ± S.E. | Mean ± S.E. | Mean ± S.E. |
| Adult Exposure | Female | 5V5M | 0 | 0 | 3 ± 0.0 | 3 ± 0.0 |
| | | 5V0G | 0 | 0 | 3 ± 0.0 | 3 ± 0.0 |
| | | 2920X | 0 | 0 | 3 ± 0.0 | 3 ± 0.0 |
| | | 5058 | 0 | 0 | 3 ± 0.0 | 3 ± 0.0 |
| | Female | 5V5M | 0 | 0 | 11 ± 0.0 | 11 ± 0.0 |
| | | 5V0G | 0 | 0 | 11 ± 0.0 | 11 ± 0.0 |
| | | 2920X | 0 | 0 | 11 ± 0.0 | 11 ± 0.0 |
| | | 5058 | 0 | 0 | 11 ± 0.0 | 11 ± 0.0 |
| | Male | 5V5M | 0 | 0 | 6 ± 0.3 | 6 ± 0.3 |
| | | 5V0G | 0 | 0 | 5 ± 0.0 | 5 ± 0.0 |
| | | 2920X | 0 | 0 | 6 ± 0.3 | 6 ± 0.3 |
| | | 5058 | 0 | 0 | 6 ± 0.3 | 6 ± 0.3 |
| Lifetime Exposure | Female | 5V5M | 3 | 4 ± 0.0 | 12 ± 0.0 | 19 ± 0.0 |
| | | 5V0G | 3 | 4 ± 0.0 | 12 ± 0.2 | 19 ± 0.2 |
| | | 2920X | 3 | 4 ± 0.2 | 12 ± 0.0 | 19 ± 0.2 |
| | | 5058 | 3 | 4 ± 0.2 | 13 ± 0.2 | 20 ± 0.0 |
| | Male | 5V5M | 3 | 4 ± 0.0 | 9 ± 0.3 | 16 ± 0.3 |
| | | 5V0G | 3 | 4 ± 0.0 | 8 ± 0.2 | 15 ± 0.2 |
| | | 2920X | 3 | 4 ± 0.1 | 9 ± 0.3 | 16 ± 0.2 |
| | | 5058 | 3 | 4 ± 0.2 | 9 ± 0.0 | 16 ± 0.2 |
| Developmental Exposure | Female | 5V0G | 3 | 4 ± 0.0 | 0 | 7 ± 0.0 |
| | | 2920X | 3 | 3 ± 0.3 | 0 | 6 ± 0.3 |
| | | 5058 | 3 | 4 ± 0.3 | 0 | 7 ± 0.3 |
| | Male | 5V0G | 3 | 4 ± 0.0 | 0 | 7 ± 0.0 |
| | | 2920X | 3 | 4 ± 0.1 | 0 | 7 ± 0.1 |
| | | 5058 | 3 | 4 ± 0.0 | 0 | 7 ± 0.0 |

**Supplemental Table 5. Sample sizes for LE and DE models.**

| Model | Sex | Diet | Total # of Cages | Total # of Mice in Each Cage | Total # of Mice |
|---|---|---|---|---|---|
| Lifetime Exposure | Female | 5V5M | 3 | 4, 3, 1 | 8 |
| | | 5V0G | 4 | 1, 1, 2, 2 | 6 |
| | | 2920X | 3 | 1, 2, 3 | 6 |
| | | 5058 | 3 | 1, 2, 3 | 6 |
| | Male | 5V5M | 4 | 3, 3, 3, 3 | 12 |
| | | 5V0G | 3 | 2, 2, 2 | 6 |
| | | 2920X | 4 | 1, 2, 2, 5 | 10 |
| | | 5058 | 3 | 1, 2, 2 | 5 |
| Developmental Exposure | Female | 5V0G-5V5M | 2 | 2, 2 | 4 |
| | | 2920X-5V5M | 2 | 1, 3 | 4 |
| | | 5058-5V5M | 2 | 2, 2 | 4 |
| | Male | 5V0G-5V5M | 2 | 2, 2 | 4 |
| | | 2920X-5V5M | 5 | 1, 1, 1, 2, 3 | 8 |
| | | 5058-5V5M | 1 | 3 | 3 |

### Figures & Figure Captions

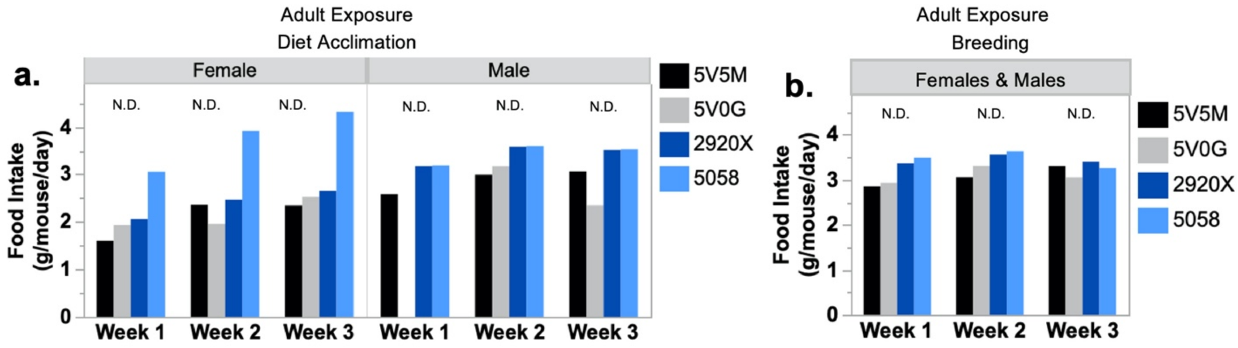


**Supplemental Figure 1. Weekly food intake measures for AE model.** Food intake was measured weekly as g/mouse/day for females and males during **a.** diet acclimation and **b.** breeding. Diet effects were calculated by one-way ANOVA or Kruskal-Wallis test and *significant when $p<0.05$. N.D. = p-value not determined due to small sample size.

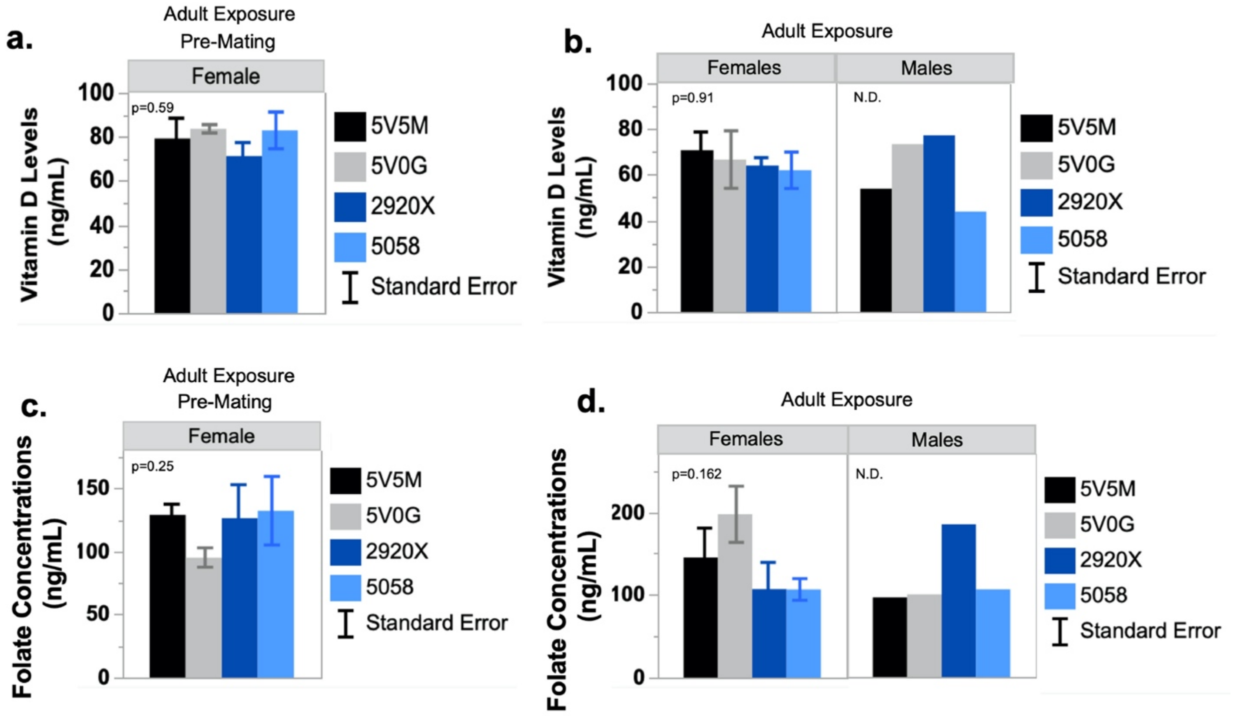


**Supplemental Figure 2. Standard chow diets do not impact serum micronutrient levels for AE model.** 25(OH)D levels were measured in serum by ELISA in **a.** females euthanized before mating (n = 5/diet) and **b.** females (n = 4/diet) and males (n = 2/diet) euthanized after mating. Folate concentrations were measured in serum using the *Lactobacillus casei* microbiological assay in **c.** females euthanized before mating (n = 5/diet) and **d.** females (n = 4/diet) and males (n = 2/diet) euthanized after mating. Error bars represent the standard error of the mean for all graphs. Diet effects were calculated by one-way ANOVA or Kruskal-Wallis test and *significant when $p<0.05$. N.D. = p-value not determined due to small sample size.

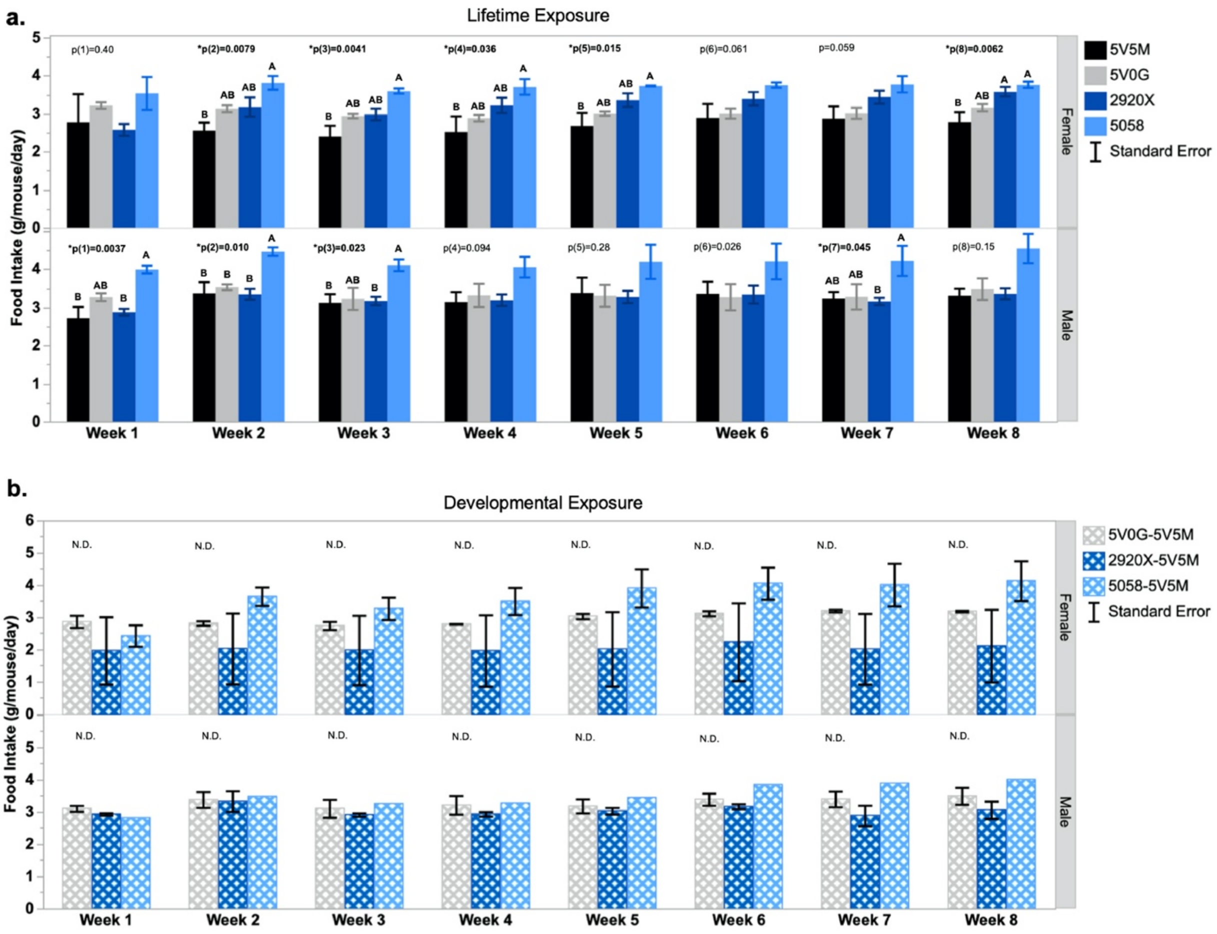


**Supplemental 3. Weekly measures of food intake for LE and DE models.** Food intake was measured weekly for each cage as g/mouse/day for females and males for **a.** LE and **b.** DE models. LE and DE models were analyzed separately. Error bars represent the standard error of the mean for all graphs. Diet effects were calculated by one-way ANOVA or Kruskal-Wallis test and *significant when $p<0.05$. N.D. = p-value not determined due to small sample size.

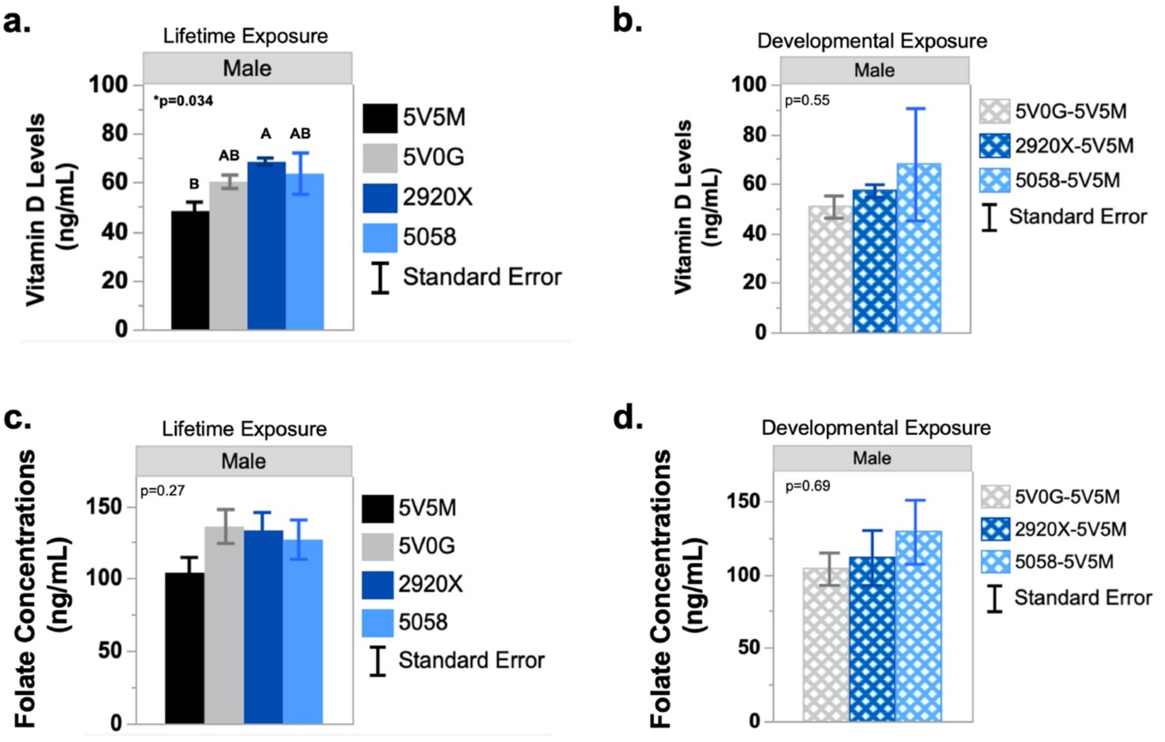


**Supplemental Figure 4. Standard chow diets impact serum vitamin D levels in LE model but not DE model and have no effect on serum folate concentrations in either model.** 25(OH)D levels were measured in serum by ELISA in **a.** LE (n = 5/diet) and **b.** DE (5V0G n = 4, 2920X n = 5, & 5058 n = 3) models. Folate concentrations were measured in serum using the *Lactobacillus casei* microbiological assay in **c.** LE (n = 5/diet) and **d.** DE (5V0G n = 3, 2920X n = 5, & 5058 n = 3) models. LE models and DE models were analyzed separately. Error bars represent the standard error of the mean for all graphs. Diet effects were calculated by one-way ANOVA or Kruskal-Wallis test and *significant when $p<0.05$.

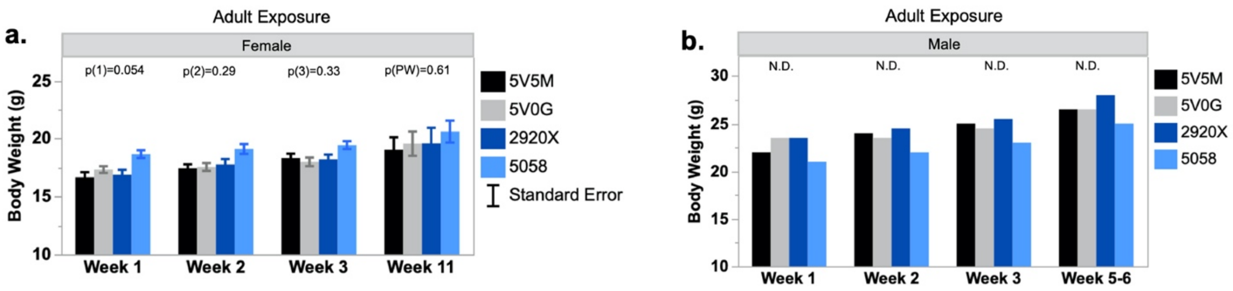


**Supplemental Figure 5. Weekly body weights for AE model.** Weekly body weights are presented for **a.** females and **b.** males. Error bars represent the standard error of the mean for all graphs. Error bars represent the standard error of the mean for all graphs. Diet effects were calculated by one-way ANOVA or Kruskal-Wallis test and *significant when $p<0.05$. N.D. = p-value not determined due to small sample size.

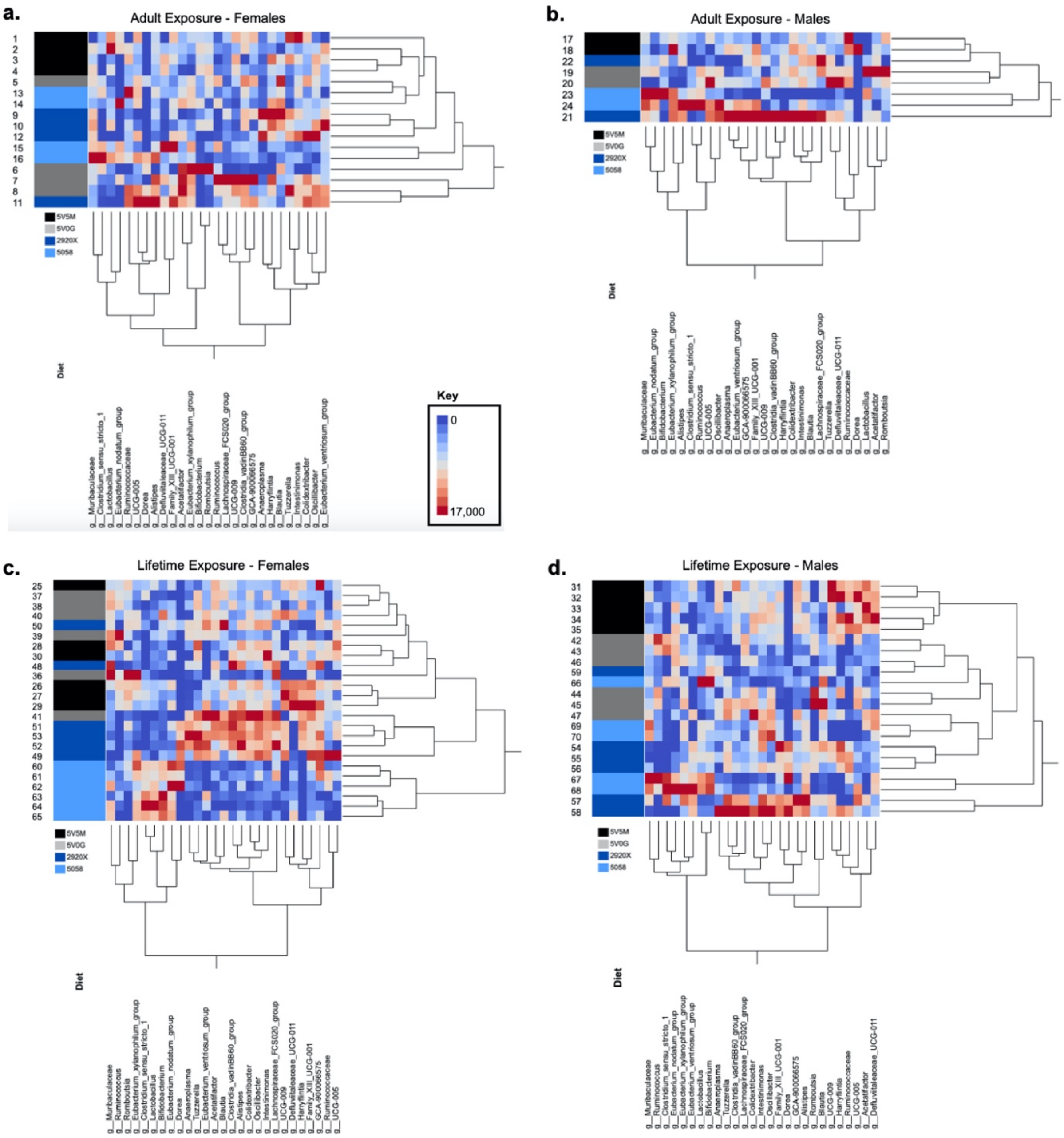


**Supplemental Figure 6. Two-way hierarchical clustering of gut bacterial genera in AE and LE models.** Heat maps of differentially abundant microbes in **a.** AE females, **b.** AE males, **c.** LE females, and **d.** LE males.
